# Age-related susceptibility to insulin resistance is due to a combination of CPT1B decline and lipid overload

**DOI:** 10.1101/2021.02.04.429529

**Authors:** Marcel A. Vieira-Lara, Marleen B. Dommerholt, Wenxuan Zhang, Maaike Blankestijn, Justina C. Wolters, Fentaw Abegaz, Albert Gerding, Ydwine van der Veen, Theo H. van Dijk, Ronald van Os, Dirk-Jan Reijngoud, Johan W. Jonker, Janine K. Kruit, Barbara M. Bakker

## Abstract

**BACKGROUND:** Advanced age increases the susceptibility to diet-induced insulin resistance (IR). A key driver of this phenomenon is lipid accumulation in the skeletal muscle. It is debated, however, whether this is due to dietary lipid overload or decline of mitochondrial function. To address the interplay of diet and age in the flexibility of muscle lipid and glucose handling, we put young and aged mice on a low- or high-fat diet (HFD).

**RESULTS:** As expected, aged mice were more susceptible to IR when given a HFD than young mice. The HFD induced intramuscular lipid accumulation specifically in aged mice, including C18:0-containing ceramides and diacylglycerols. This was reflected by the mitochondrial β-oxidation capacity, which was upregulated by the HFD in young, but not in old mice. Conspicuously, most β-oxidation proteins were upregulated by the HFD in both groups, but carnitine palmitoyltransferase 1B (CPT1B) declined in aged animals. Computational modelling traced the flux control mostly to CPT1B, suggesting a CPT1B-driven loss of flexibility to the HFD with age. Finally, in old animals glycolytic protein levels were reduced and less flexible to the diet.

**CONCLUSION:** We conclude that intramuscular lipid accumulation and decreased insulin sensitivity are not due to age-related mitochondrial dysfunction or nutritional overload alone, but rather to their interaction. Moreover, we identify CPT1B as a potential target to counteract age-dependent intramuscular lipid accumulation and thereby IR.

## Background

The skeletal muscle is one of the principal tissues to increase postprandial glucose uptake in response to insulin. If the normal metabolic response to insulin is compromised, a condition known as insulin resistance (IR) arises (1). Given its major role in maintaining whole-body glucose homeostasis, skeletal muscle IR precedes the onset of type 2 diabetes (2). Lipid-induced IR is the most important form of IR in the skeletal muscle (1, 3, 4), for which ageing is a risk factor (5).

Different mechanisms have been proposed to explain how lipids could interfere with normal glucose uptake and utilization in the skeletal muscle. The first mechanism, initially described by Randle, is a fatty-acid-glucose cycle in which fatty acids and glucose reciprocally inhibit each other’s oxidation by allosteric enzyme inhibition (6). Since then, the focus has shifted to the insulin signalling cascade, which triggers recruitment of the glucose transporter GLUT4 to the plasma membrane (1). Diacylglycerols (DG’s) and ceramides (Cer’s) are the two major acknowledged lipid classes to cause IR by inhibiting the insulin signalling cascade (1, 4, 7).

The accumulation of such lipid species and lipid-induced IR can be ascribed to an imbalance between fatty-acid uptake and mitochondrial oxidation (7-9). The current debate concerns the question of whether lipids accumulate due to mitochondrial dysfunction (9) or as a result of an overload by excess nutrients (10, 11). On the one hand, the idea of an underlying mitochondrial defect is supported by reports that increasing the fatty-acid β-oxidation capacity protects against IR (12, 13). Moreover, increased age has been associated with a decrease in mitochondrial function across different species (14-16). On the other hand, different studies have shown that inhibiting the first steps of β-oxidation can prevent the development of lipid-induced IR (10, 17, 18), which led to the idea that overloading the pathway causes a metabolic ‘gridlock’ (11). In line with this, computer simulations suggested that the β-oxidation is intrinsically susceptible to such a gridlock by an accumulation of intermediate metabolites, leading to inhibition of downstream metabolism (19).

In this study we address the interaction between mitochondrial dysfunction and nutritional overload in mice. A potential interaction between both mechanisms is supported by the finding that advanced age leads to a higher susceptibility to high-fat-diet (HFD)-induced IR due to changes in fatty-acid handling (20, 21). We investigated by lipidomics profiling how advanced age exacerbates HFD-induced lipid accumulation in mouse quadriceps and its relation to peripheral insulin sensitivity. Subsequently, we used a systems-biology approach to characterize mitochondrial lipid handling by high-resolution respirometry, targeted proteomics, and computational modelling. We found that aged mice lose their mitochondrial flexibility in the quadriceps to respond to a HFD and identified a specific role for CPT1B.

### Research Design & Methods

More detailed description can be found online in the Supplemental Material.

### Animals & Diet

Male C57BL/6J mice were obtained from the Mouse Clinic for Cancer and Aging of the University of Groningen, and kept under standard housing conditions with *ad libitum* access to food (rodent chow diet (RM1) SDS Diets, Woerden, The Netherlands) and water, a 12h light/dark cycle and a temperature-controlled environment. Before being fed experimental diets, 3 and 18-month old mice (referred to as young and old, respectively) were fed a semi-synthetic LFD and were assigned to either a 20% low-(LFD) or 60% high-fat diet (HFD). (based on the AIN-93G breeding diet (D10012G), Open Source Diets, New Brunswick, NJ, USA, Supp. Table 1) for 12 weeks. Prior to the experiment, mice were put on a 2-week run-in period, containing a control diet (10% fat) to normalize microbial health. One week before experimental diets were introduced, mice were housed individually. Supp. Fig. 1A: detailed timeline. Energy expenditure (EE), respiratory exchange ratio (RER) and measurements from blood samples were performed as described elsewhere (22).

### Assessment of glucose and insulin tolerance

To determine changes in glucose homeostasis and insulin sensitivity, an oral glucose tolerance test (OGTT) was performed following oral administration of 1.5g/kg body weight D-glucose after overnight (10h) fasting. A 25% glucose solution (w/v) was given by oral gavage. Glucose was measured using a OneTouch Select Plus glucose meter (Lifescan, Zug, Switzerland). Insulin concentrations were determined using the rat insulin ELISA kit from Crystal Chem (Cat. 90010, Zaandam, The Netherlands) and a mouse insulin standard (Cat. 90020, Zaandam, The Netherlands). Homeostatic Model Assessment of Insulin Resistance (HOMA-IR) indexes were calculated as previously described using both the fasting insulin and fasting glucose levels and corrected for mice (23). The Muscle Insulin Sensitivity Index (MISI) calculation was adapted from (24).

### Lipid extraction, LC-MS analysis and data processing

Frozen quadriceps tissue was homogenized in 0.9% NaCl (15% w/v) using a BeadBeater system (Precellys^®^ Evolution, Bertin Technologies). Lipid extraction was performed following the protocol of Matyash *et al*. (25) with slight modifications as described in the supplemental materials and methods. LC-MS lipid analysis was performed on an Ultimate 3000 High-Performance UPLC coupled with a Q Exactive MS (Thermo Fischer Scientific, Darmstadt, Germany). Data acquisition was performed with Thermo Xcalibur® software [(version 3.2.63), Thermo Scientific, Waltham, MA], Progenesis QI® software (Waters Corporation, Milford, MA) and Lipidhunter2 software (26).

### Mitochondrial isolation and respiration

Mitochondria were isolated from fresh quadriceps as previously described with adaptations (16). Mitochondrial suspensions were incubated at 37 °C in a two-channel high-resolution Oroboros oxygraph-2k (Oroboros, Innsbruck, Austria) according to supplement materials. Malate (2 mM) was present in all assays. 2 mM pyruvate (with or without 5 mM glutamate) or 25 µM palmitoyl-CoA, the latter supplemented with 2 mM carnitine, were used as substrates.

### Proteomics

15% (w/v) quadriceps homogenates were prepared in DPBS (Gibco 14190) with a BeadBeater system (Precellys^®^ Evolution, Bertin Technologies). Samples were centrifuged at 15,000 *g* for 5 minutes, supernatants were collected and cOmplete(tm) proteinase inhibitor cocktail was added (1:25, Merck 11836145001). Protein concentrations were measured with the Pierce BCA Protein Assay Kit (ThermoFisher 23225). Isotopically-labelled standard peptides for mitochondrial targets had been previously selected (27) and additional peptides were selected for enzymes in glucose metabolism (table with sequences: Supp. Table 7). In-gel digestion with trypsin, LC-MS analysis and data analysis have been performed according to Wolters et al. (27).

### Enzyme activities

Hexokinase and pyruvate kinase activities were carried out using NAD(P)H-linked assays. Citrate synthase activity was measured by dithionitrobenzoate (DTNB) oxidation. Both were performed at 37 °C in a Synergy H4 plate reader (BioTek(tm)) at 340 nm and 412 nm, respectively. Carnitine palmitoyltransferase (CPT) activity was measured by following the production of palmitoylcarnitine with LC/MS-MS.

### Western Blots

For Western Blot analysis, quadriceps homogenates (7.5% w/v) in NP-40 buffer were used. Electrophoresis, transfer, antibody incubations and detection were performed as previously reported (28).

### Computational modelling

The computational model of mouse mitochondrial β-oxidation previously described (29) was converted to .nb format (Wolfram Mathematica, Wolfram Research, Inc., Champaign, IL, USA). A full description of the modelling strategy and the script can be found in Supplemental text 1 and Appendix 1, respectively.

### Statistical analysis

Data were visualized using GraphPad Prism software (GraphPad Software Inc., version 8.0, 2018) and analysed with either the same software or IBM SPSS Statistics (IBM Corp., Version 25.0, 2017). Data are expressed as mean ± SEM unless otherwise stated. Analyses were conducted using Student’s t-test, 2- and 3-way ANOVA.

## Results

### Ageing increases susceptibility to develop diet-induced insulin resistance

Young (3 months) and old (18 months) C57BL/6J mice were given either a low-(LFD) or high-fat (HFD) for 12 weeks. The diets consisted of 20% and 60% of kcal from fat, respectively (Supp. Table 1). Despite old mice being already heavier than young mice, both age groups fed with a HFD gained more weight than their LFD counterparts (Fig. 1A-B). Young and old mice gained the same absolute quantity of body fat (Fig. 1C) and body weight (Supp. Fig. 1B) on the HFD. Aged mice had, on average, a 17% higher food intake than young mice, in agreement with higher body weight (Supp. Fig 1B). Respiratory exchange ratios (RER) showed a substrate switch from carbohydrate (high RER) to fat (low RER) in response to the HFD (p_diet<_0.0001) (Fig. 1D; Supp. Fig. 1D). After adjustment for body weight, higher energy expenditure was observed in mice fed a HFD versus LFD independently of age (p_diet_<0.0001, Fig. 1E). Mouse physical activity did not differ between age groups, but was decreased by the HFD (p_diet_<0.0001; Supp. Fig. 1E). In the oral glucose tolerance test (OGTT), HFD-treated groups were more glucose intolerant than LFD groups (Fig. 1F-G), while plasma insulin levels were much higher in old animals than in young animals, reaching around 5 ng/mL in HFD-old mice (Fig. 1H-I). From fasting insulin and glucose levels we calculated the HOMA-IR index as a parameter for total body IR. The HOMA-IR was strongly increased by both age and diet (p_age_=0.0002, p_diet_<0.0001). Interestingly, the HOMA-IR of old mice increased more strongly with the HFD than that of young mice (Fig. 1J). The Muscle Insulin Sensitivity Index (MISI) estimates the rate of glucose removed per minute over average insulin concentration during the OGTT (24). Based on this parameter, the HFD acted on both age groups to lower the insulin sensitivity, which was further decreased in old animals (p_age_ and p_diet_<0.0001) (Fig. 1K). Surprisingly, old mice had decreased levels of non-esterified fatty acids (NEFAs, p_age_=0.007) when compared to young animals (Fig. 1L). Plasma TGs were also decreased by advanced age to a higher extent and an increase of about 30% was observed in both age groups when fed a HFD (p_diet_=0.009) (Fig. 1M). No differences were observed in the plasma levels of branched-chain amino acids, previously associated with the development of IR (30) (Supp. Fig 1F). Altogether, these data show that HFD-induced insulin resistance was exacerbated in the aged animals.

**Figure 1:**
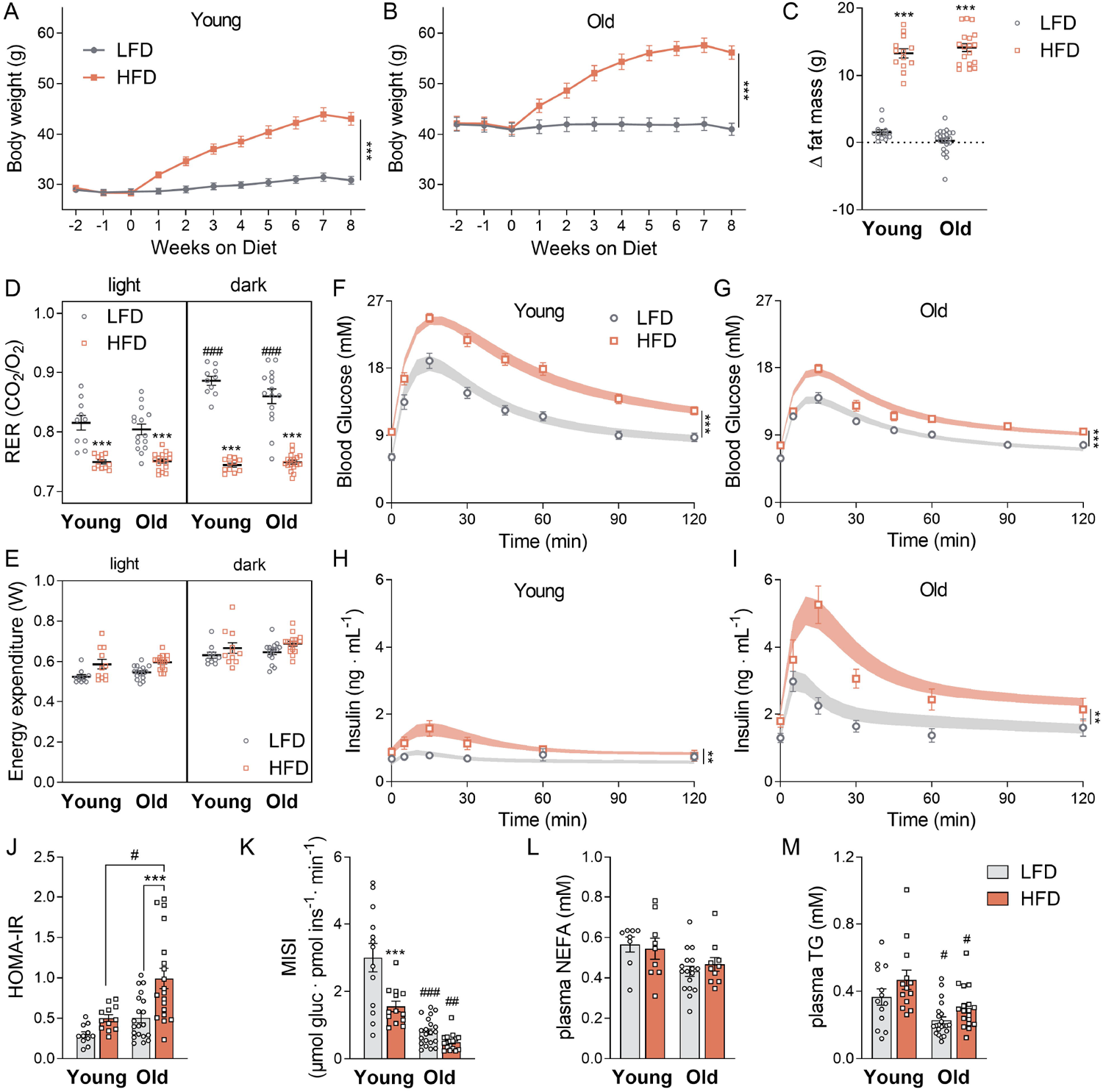
Aged mice are more susceptible to develop diet-induced insulin resistance. A-B: Weight increase during 8 weeks in young and old mice, respectively, when submitted to either a LFD or HFD. C: Increase in fat mass after 8 weeks on a HFD. D: Average respiratory exchange ratios (RER) in both light/dark phases following 11 weeks on each diet. E: Energy expenditure in both light/dark phases after 11 weeks on each diet. Data were corrected with ANCOVA with BW as covariate for both light phases in old and aged mice (adjusted body weight = 45 g). Plasma glucose concentrations for young (F) and old (G) mice and plasma insulin levels for young (H) and old (I) mice are plotted for 120 min following an oral glucose tolerance test (OGTT) (1.5g/kg BW glucose), performed after 9 weeks on either a LFD or a HFD. Points represent experimental data and fitted curves are summarized in the shaded area. J: HOMA-IR indexes adjusted to mice were calculated from the OGTT data as [fasting glucose]×[insulin]/14.1. K: Muscle Insulin Sensitivity Index (MISI) calculated from fitted glucose and insulin curves, adapted from O’Donovan et. al, 2019 (24). L-M: plasma non-esterified fatty acids (NEFAs) and triglycerides (TG). Data are shown as mean ± SEM. Statistical analysis was conducted for each group according to the Materials & Methods section, n=13-22 per group, ^###^p<0.001 (old vs young), ^##^p<0.01 (old vs young), ^#^p <0.05 (old vs young), ***p<0.001 (LF vs HF), **p<0.01 (LF vs HF), *p<0.05 (LF vs HF). Each pairwise comparison analyses means that differ by only one factor, meaning that age comparisons have matched diet and diet comparisons have matched age.

### Extensive lipidome remodelling in the quadriceps of old, but not young mice on a HFD

To test whether the susceptibility of aged mice to develop IR is associated with changes in the skeletal muscle lipidome, we used an untargeted lipidomics approach. In total we detected 443 lipid species in the quadriceps samples (Fig. 2A; Supp. Table 2). In young mice the HFD caused only minor changes in the lipidome, with 7 lipid species mildly increased and 2 lipid species decreased (Fig. 2B and Supp. Table 3). In contrast, in old mice, a substantial number of 58 lipid species were higher in the HFD group than in the LFD group, and 3 lipid species were decreased (Fig. 2C and Supp. Table 3). A lipid class that was specifically upregulated in old mice on HFD comprised long-chain acylcarnitines (Fig. 2D). This was confirmed by targeted LC-MS for C18:0-carnitine (Fig. 2E, complete profiling: Supp. Table 4). Such accumulation of long-chain acylcarnitines could represent an imbalance in old mice between uptake and consumption of fatty acids. This could then drive the conversion into complex lipid species that cause IR. Cer(d18:1/18:0) was previously demonstrated to induce IR, specifically in muscle (31). This lipid was increased by the HFD in our study in an age-dependent manner (Fig. 2F). Overall, C18:0 side chains were overrepresented in the HFD-induced lipids (Supp. Table 3). For instance the diacylglycerol DG(16:0_18:0) was induced by the HFD in old mice (Fig. 2G). Finally, out of the 11 TGs increased by the HFD, all were increased only in old mice and TG(18:0_18:0_18:1) showed the highest fold change (7.7; Fig. 2H).

**Figure 2:**
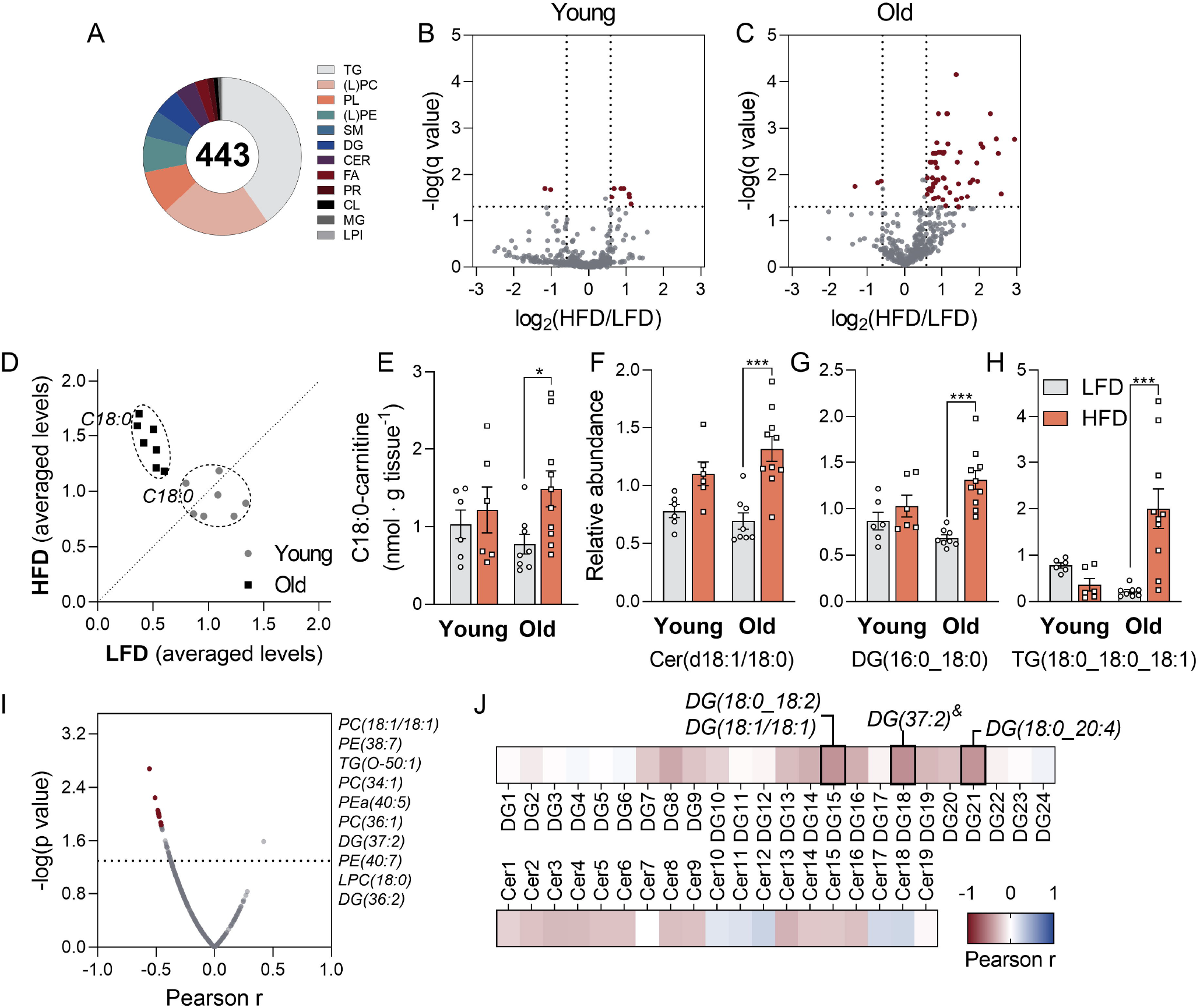
HFD causes extensive lipid remodelling in the quadriceps of aged mice. A: Number of detected lipid species per category. TG: triglycerides, (L)PC: (lyso)phosphatidylcholine, PL: plasmalogen, (L)PE: (lyso)phosphatidylethanolamine, SM: sphingomyelin, DG: diacylglycerol, CER: ceramides, FA: fatty acids, PR: prenol lipids, CL: cardiolipin, MG: monoacylglycerols, LPI: lysophosphatidylinositol. Volcano plots comparing the effect of a HFD (vs LFD) on young (B) and old (C) mice, corrected for multiple comparisons with the False Discovery Rate (FDR) method, with Q = 5%. A fold change threshold was defined as higher than 1.5 or smaller than 0.66. D: Scatter plot of normalized levels of long-chain acylcarnitines detected via untargeted lipidomics (C14:0, C16:0, C18:0, C20:0, C16:0-OH, C18:0-OH, C18:1-OH). Each dot represents the mean value of a specific metabolite on the HFD (y-axis) versus the LFD (x-axis). Prior to plotting, the levels of each acylcarnitine were normalized to the average value of the same acylcarnitine in all animals. Metabolites along the dashed line are not regulated by the diet (mean LFD = mean HFD), while metabolites above or below the line are up- or downregulated, respectively, by the HFD. E: Concentrations of C18:0 acylcarnitines detected by LC-MS. F-H: Comparison between diets and age for Cer(d18:1/18:0), DG(16:0_18:0) and TG(18:0_18:0_18:1), respectively. I: Pearson correlation coefficients (r) between detected lipid species and MISI (x-axis) plotted against -log(p-value) (y-axis). Top 10 lipids with highest correlation coefficients are specified next to the figure. J: Heatmap for Pearson correlation coefficients for detected DG’s and Cer’s (from all groups) versus MISI (&low signal lipid, only summed structure information). Statistically significant correlations (p<0.05) are highlighted. Data are shown as mean ± SEM, n=6-10. Statistical analysis was conducted for each group according to the Materials & Methods section, ^###^p<0.001 (old vs young), ^##^p<0.01 (old vs young), ^#^p <0.05 (old vs young), ***p<0.001 (LFD vs HFD), **p<0.01 (LFD vs HFD), *p<0.05 (LFD vs HFD).

Among the detected lipids, 39 species correlated negatively with the MISI index with p<0.05 (Fig. 1K, 2I). PC(18:1/18:1) had the strongest correlation (r = −0.56, p=0.002, Supp. Fig. 2). A direct role of this phosphatidylcholine, however, has not been previously addressed. Given the previously established roles of ceramides (Cer’s) and diacylglycerols (DG’s) on the insulin signalling pathway (1, 4), we focused on the association between the detected Cer’s and DGs and the MISI index. We found three DG’s and no Cer significantly negatively associated with MISI (Fig. 2J, Supp. Table 5).

In conclusion, we found that old mice were more prone than young mice to accumulate lipids in the quadriceps in response to a HFD, particularly C18:0-containing species, and that this correlated with the MISI index.

### Mouse quadriceps display decreased mitochondrial flexibility during ageing

The observed lipid accumulation in old mice on a HFD suggests that oxidation cannot meet the increased lipid supply in these animals. To assess the muscle oxidative capacity, we analysed mitochondrial content and function. The protein levels of the peroxisome proliferator-activated receptor γ coactivator 1-alpha (PGC-1α), a regulator of mitochondrial biogenesis, were unchanged among the groups (Fig. 3A). Citrate synthase activity in cell homogenates (CS), a marker of mitochondrial mass, was not changed by age and hardly affected by the diet (Fig. 3B). Also in isolated mitochondria the CS activity remained unaltered (Fig. 3B). O_2_ consumption was measured in isolated mitochondria (Table 1) and then expressed per total tissue protein (Fig. 3C-E). Young mice had an 83% higher oxidative capacity towards palmitoyl-CoA on a HFD than on a LFD (p<0.01). In contrast, the HFD did not significantly affect oxidative capacity in old mice (Fig. 3C). Diet and age did not affect O_2_ consumption when pyruvate or pyruvate with glutamate were used as substrates (Fig. 3D-E). Together, these results suggest specific alterations in the β-oxidation pathway, rather than in the TCA cycle or electron transport chain (ETC), since the latter are shared by all three substrates. Apparently, ageing does not cause mitochondrial deficiency *per se*, but instead blunts the flexibility of the β-oxidation pathway to respond to the different diets.

**Table 1:** O_2_ consumption rates (States 3 and 4) for isolated mitochondria from quadriceps. All substrates were added in the presence of malate. Values are shown as mean ± SEM (nmol · mg mitochondrial protein^-1^ · min^-1^). State 3: ADP-stimulated maximal respiration. State 4: carboxyatractyloside-induced inhibition of ATP synthesis (via ATP-ADP translocase inhibition), *p_diet_=0.025

|  |  | Substrates |  |  |  |  |  |
| --- | --- | --- | --- | --- | --- | --- | --- |
| Tissue |  | pyruvate |  | pyruvate/glutamate |  | palmitoyl-CoA/<br>carnitine |  |
| Quadriceps |  | YOUNG | OLD | YOUNG | OLD | YOUNG | OLD |
| State 3 | LFD | 428± 56 | 424± 58 | 476 ± 68 | 458 ± 57 | 131± 29 | 153 ± 20 |
|  | HFD | 524 ± 56 | 415± 48 | 627 ± 77 | 419 ± 43 | 204 ± 14* | 184 ± 20* |
| State 4 | LFD | 20.9 ± 5.9 | 20.2 ± 4.5 | 21.9 ± 6.8 | 30.2 ± 9.7 | 36.1 ± 10.2 | 26.5 ± 3.6 |
|  | HFD | 20.2 ± 2.6 | 18.9 ± 3.7 | 27.6 ± 8.1 | 14.9 ± 2.2 | 35.6 ± 4.4 | 26.1 ± 5.1 |
| RCR | LFD | 26.0 ± 4.7 | 26.3 ± 3.7 | 38.6 ± 13.1 | 25.3 ± 3.9 | 4.2 ± 0.8 | 6.1 ± 0.5 |
|  | HFD | 28.2 ± 5.1 | 27.0 ± 4.0 | 30.3 ± 10.9 | 31.8 ± 3.9 | 5.9 ± 0.5 | 7.6 ± 0.8 |

**Figure 3:**
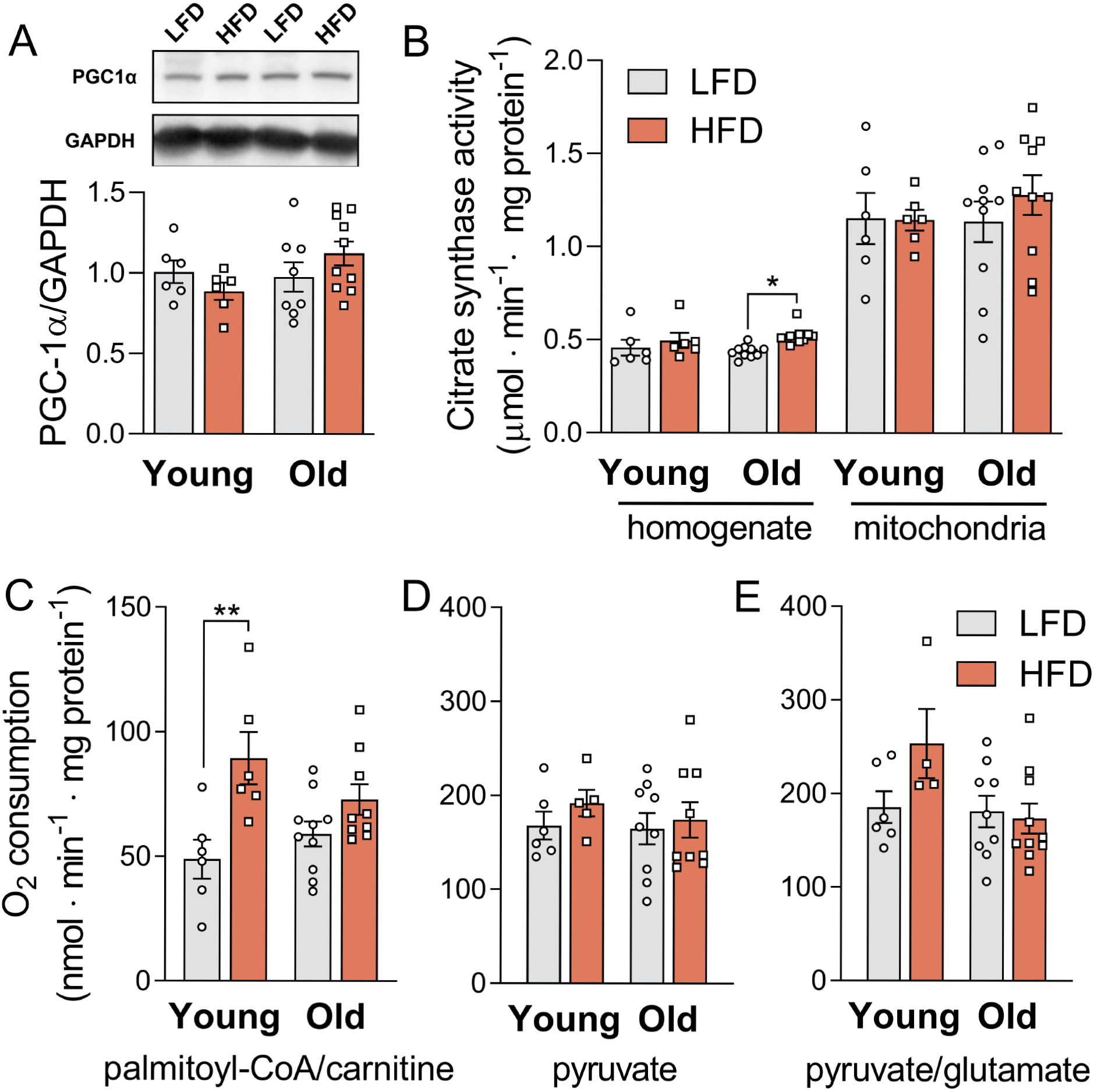
A HFD increases mitochondrial capacity to oxidize fatty-acids only in young mice. A: Total PGC1α protein content in quadriceps homogenates normalized by glyceraldehyde 3-phosphate dehydrogenase (GAPDH) levels. B: Citrate synthase activities measured in quadriceps homogenates and mitochondria enriched suspensions. C: Scheme with respiratory substrates used for experiments. Mitochondrial O_2_ consumption fluxes were corrected for mitochondria enrichment of suspensions using the ratio between citrate synthase activity (CS) in the mitochondria versus in total homogenate (R=CS_mito_/CS_homog_). Maximal ADP-stimulated O_2_ consumption per tissue protein is shown for palmitoyl-CoA and carnitine (D), pyruvate (E) and pyruvate and glutamate (F) as substrates, all in the presence of malate. Data are shown as mean ± SEM and were analysed using Two-Way ANOVA. Multiple comparisons were performed using Tukey’s correction. N=6-10 per group, **p<0.01 (HFD vs LFD), *p<0.05 (HFD vs LFD), matched ages.

### Ageing downregulates CPT1B despite HFD-induced β-oxidation remodelling

To explain the loss of flexibility of mitochondrial β-oxidation, we took a targeted and quantitative proteomics approach, based on isotopically-labelled peptide standards. 16 mitochondrial β-oxidation proteins were quantified, 14 of which were increased by the HFD relative to the LFD in both age groups (p_age_<0.05) (Fig. 4A and Supp. Table 6). Proteins clustered together per age group, with the highest protein concentrations in old mice (Fig. 4A). The exception was carnitine palmitoyltransferase 1B (CPT1B, muscle isoform); both protein levels and activity were increased by the HFD, but decreased with advanced age (p_age_ and p_diet_ < 0.05) (Fig. 4B-C). Closer inspection of both CPT1B protein concentration and enzyme activity showed a similar loss of flexibility that was earlier observed in the lipidome: in young mice on HFD protein and activity reached levels around 90% and 50% higher than in old mice, respectively (Fig. 4B and C). This loss of flexibility in old mice was also observed for very-long-chain acyl-CoA dehydrogenase (VLCAD) and medium-chain ketoacyl-CoA thiolase (MCKAT), two other key β-oxidation enzymes (19, 32) that were increased by the HFD mostly in younger animals (Fig. 4D-E).

**Figure 4:**
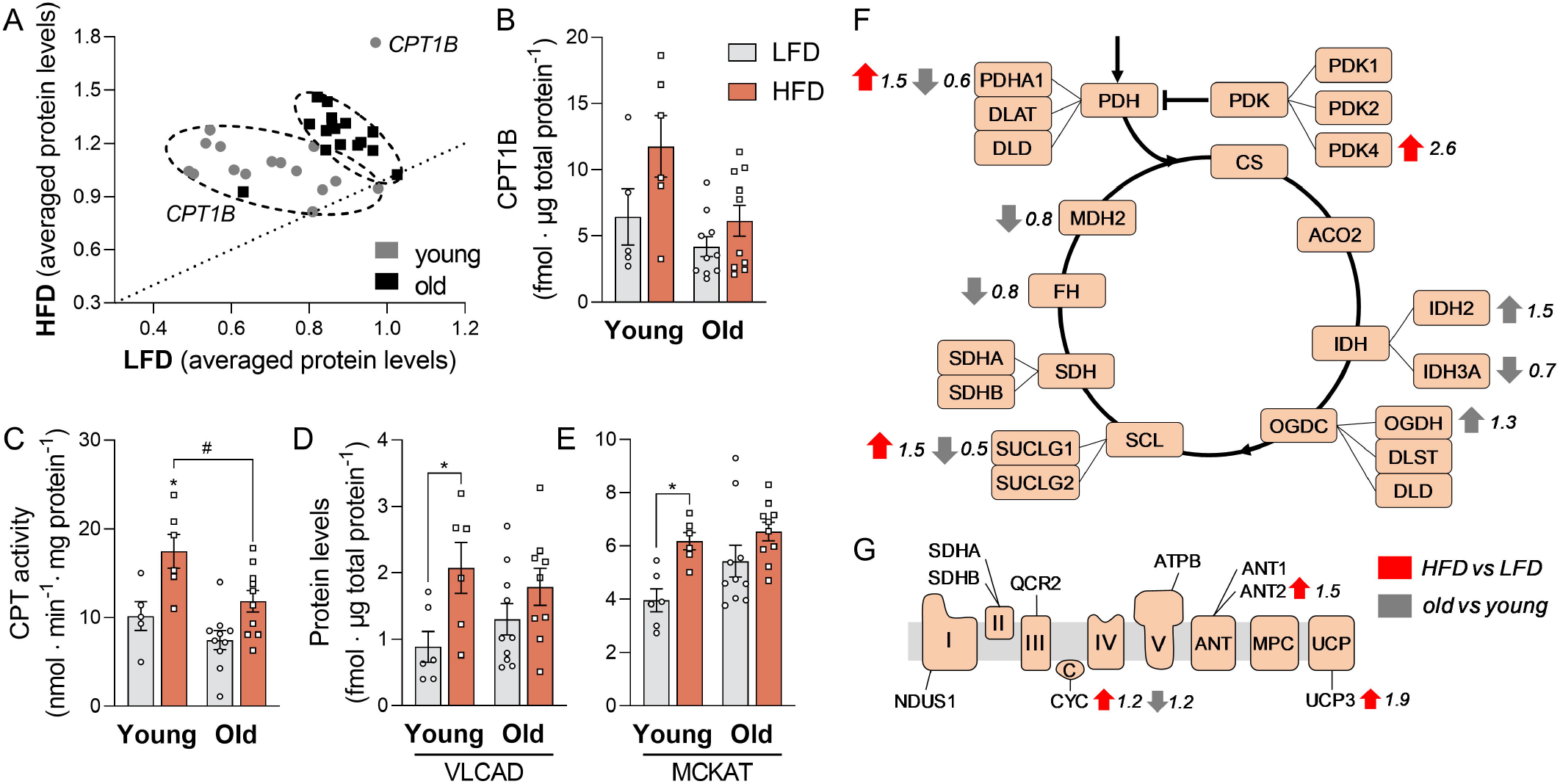
Upregulation of β-oxidation proteins by the HFD, despite accompanying loss of CPT1B with ageing. A: Scatter plot of normalized levels of β-oxidation proteins detected in mouse quadriceps via targeted proteomics Each dot represents the mean value of a specific protein on the HFD (y-axis) versus the LFD (x-axis). Prior to plotting, the levels of each protein were normalized to the average value of the same protein in all animals. Proteins along the dashed line are not regulated by the diet (mean LFD = mean HFD), while proteins above or below the line are up- or downregulated, respectively, by the HFD. CPT1B: carnitine palmitoyltransferase 1B. B: Relative CPT1B content in all groups. C: Total CPT activity measured in mitochondrial samples and corrected for the enrichment of the preparation (CS_mito_/CS_homog_) as mentioned for figure 3D. D-E: Quantification of very-long-chain acyl-CoA dehydrogenase (VLCAD) and medium-chain ketoacyl-CoA thiolase (MCKAT). F-G: Schematic representation of the effects of age and diet on the TCA cycle and ETC, respectively. Each box represents a protein that has been measured by our method and the specific isoforms or subunits that have been measured are annotated. Red arrows represent the effect of diet (HFD vs LFD) while the grey arrows indicate the age effect (old vs young) based on significant 2-way ANOVA results. PDH: pyruvate dehydrogenase complex, CS: citrate synthase, ACO2: aconitase, IDH: isocitrate dehydrogenase, OGCD: α-ketoglutarate dehydrogenase complex, SCL: succinyl-CoA ligase, SDH: succinate dehydrogenase (complex II), FH: fumarate dehydrogenase, MDH2: malate dehydrogenase, ANT: adenine nucleotide translocator, MPC: mitochondrial phosphate carrier protein, UCP: uncoupling protein. Data are shown as mean ± SEM and were analysed using Two-Way ANOVA and multiple comparisons were performed using Tukey’s correction. N=6-10 per group, **p<0.01 (HFD vs LFD), *p<0.05 (HFD vs LFD), matched ages and ^#^p <0.05 (old vs young), matched diets.

Further analysis of the downstream pathways revealed minor changes in the TCA cycle and ETC (Fig. 4F-G) by age. Pyruvate dehydrogenase kinase 4 was increased by HFD (PDK4, FC: 2.6), as reported previously (33). Increased PDK4 expression leads to inhibition of pyruvate dehydrogenase (PDH) by phosphorylation of the PDH-E1α subunit (33) and contributes to the shift from carbohydrate to fatty-acid utilization. Here the effect was, however, counteracted by an upregulation of the PDH-E1α (PDHA1, FC: 1.5) and the net effect of both changes is unclear. Uncoupling protein 3 (UCP3) was upregulated by the diet (FC: 1.9), also a known effect of increased fatty-acid availability (34). Apart from these changes, these results corroborate the notion that β-oxidation is the main pathway in energy metabolism that is affected by the HFD (35). In conclusion, the decrease in flexibility of the mitochondrial β-oxidation pathway could be traced down to a loss of flexibility and content of CPT1B with age in mice fed a HFD, with possibly a contribution by VLCAD and MCKAT.

### HFD-proteome remodelling is overruled by low and less flexible CPT1B in aged animals

While there was a substantial upregulation of most fatty-acid oxidation enzymes in both HFD-treated groups, CPT1B content and activity were significantly lower in old than in young animals. This raised the question of what is the net *in vivo* effect of these opposing responses in the proteome. We tested this computationally, using a dynamic β-oxidation model previously developed by our group (29). To parameterize it to skeletal-muscle mitochondria, the proteomics data were used (see Supp. Text 1 for details). A version of the β-oxidation model was created for each individual mouse in the dataset. The steady-state palmitoyl-CoA consumption flux, representing the conversion of palmitoyl-CoA to acetyl-CoA, was plotted as a measure of the β-oxidation. Palmitoyl-CoA concentrations were chosen within the likely physiological range of skeletal-muscle tissue (36) (Fig. 5B). The results indicated a significant diet and age effect (p_age_<0.05, p_diet_<0.05), and a stronger upregulation of the β-oxidation fluxes by the HFD in young mice, reaching values 75% higher than in aged mice in the given substrate range. *In vivo*, however, CPT1B is inhibited by endogenous malonyl-CoA (32). Acetyl-CoA carboxylase (ACC) levels, as well as its active phosphorylated form (pACC) which synthesizes malonyl-CoA, remained unchanged in all four groups (Supp. Fig. 3A-C). Malonyl-CoA decarboxylase (MCD) levels did not differ either (Supp. Fig. 3D). Therefore, we applied the same malonyl-CoA concentration to all groups. As expected, the addition of malonyl-CoA reduced the β-oxidation fluxes in the simulations (Fig. 5C), but resulted in a similar pattern among groups as observed without malonyl-CoA (Fig. 5B). Flux control coefficients (FCC) quantify how the pathway flux is modulated when the maximal capacity of a particular enzyme is changed (37), i.e. to which extent the enzyme controls the flux. In the absence of malonyl-CoA, the FCC_CPT1B_ remained approximately constant (∼1) for old animals, implying that this was nearly the sole rate-limiting step within the analysed concentration range of palmitoyl-CoA. In contrast, FCC_CPT1B_ tended to decrease with a palmitoyl-CoA overload in young animals (Fig. 5D). This drop was compensated by increased control in MCKAT (Fig. 5E). The control of CPT1B is, however, expected to remain close to 1 within a wider range of palmitoyl-CoA *in vivo* due to the action of malonyl-CoA (32), which was confirmed in our simulations (Fig. 5F). These results suggest that, despite the remodelling of most β-oxidation proteins by the HFD in both age groups, the lack of flexibility at the level of CPT1B overruled this regulation and constrained the fluxes through the pathway specifically in aged animals.

**Figure 5:**
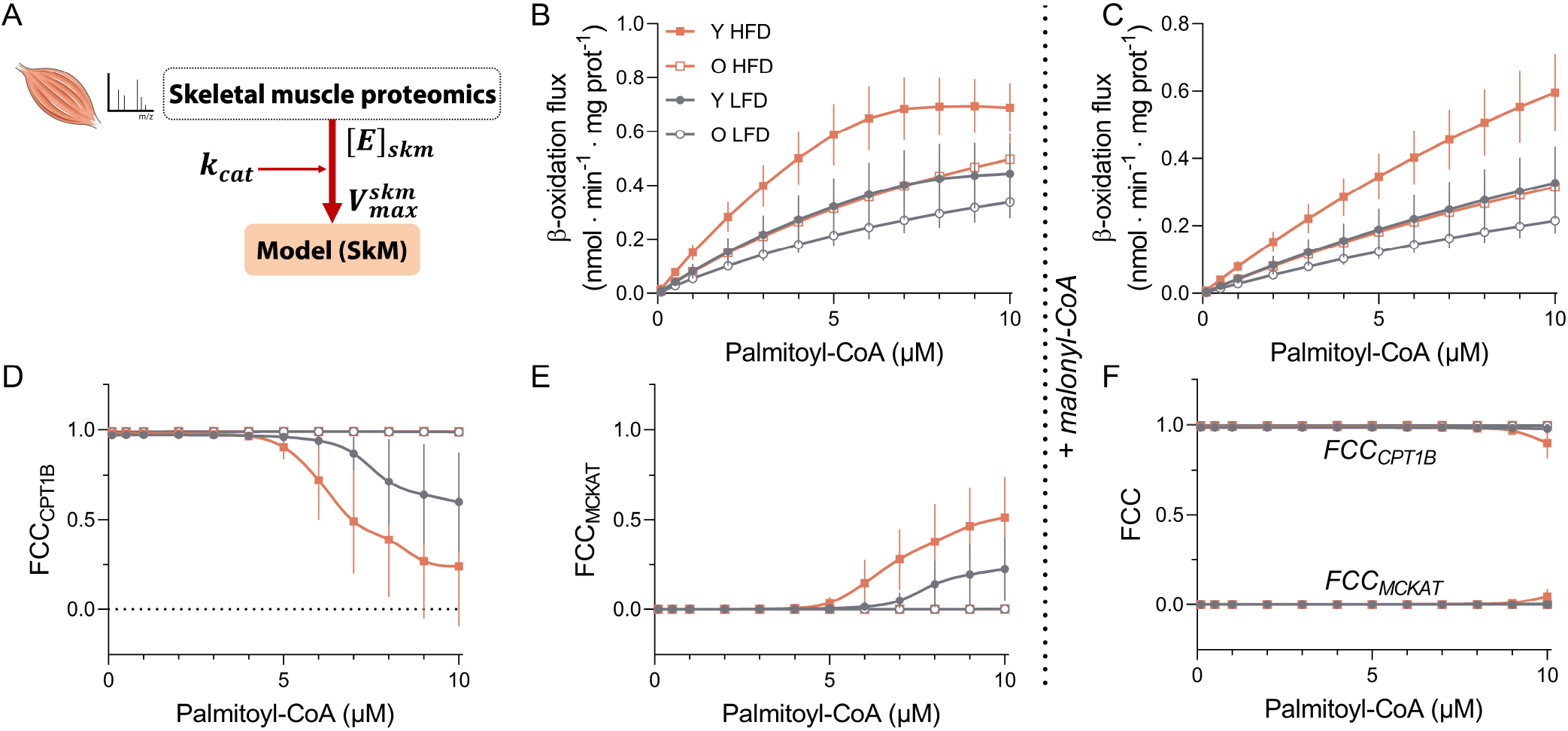
*In silico* β-oxidation simulations confirm flux dependence on CPT1B. A: Schematic representation of the strategy used to parameterize the computational model of mitochondrial β-oxidation to skeletal muscle. Proteomics data from liver previously published were used to estimate k_cat_’s for the proteins involved in the pathway and applied to proteomics data obtained from quadriceps samples to calculate V_max_’s for each animal. B: Estimated steady-state β-oxidation fluxes (defined as the fluxes through CPT1B) based on *in vivo*-like conditions for different concentrations of substrate (palmitoyl-CoA) ranging from 0.1 to 10 µM and normalized for total homogenate protein. C: Estimated steady-state β-oxidation-fluxes in the presence of 0.2 µM malonyl-CoA. Flux control coefficients (FCC) were calculated for the same substrate range. FCC is defined as *∂*J_ss_/*∂*E_i_, which represents to what extent an enzyme ‘i’ (E_i_) can alter the steady-state flux of a pathway (J_ss_). FCC’s for CPT1B (D) and MCKAT (E) are depicted. F: FCC’s for CPT1B and MCKAT in the presence of 0.2 µM malonyl-CoA. All simulations were performed for the V_max_’s estimated for each mouse in the dataset. Y stands for young (LFD: grey full circles, HFD: orange full squares) and O for old (LFD: grey open circles, HFD: orange open squares). Data are shown as mean ± SEM and were analysed using Two-Way ANOVA. A complete description of the model and the assumptions made can be found in Supp. Text 1.

### The glycolytic proteome shows less flexibility in aged mice

Finally, we investigated the flexibility of glucose metabolism at the proteome level. Again using isotopically-labelled standards peptides, we quantified 40 peptides, spanning 31 proteins involved in glycolysis and glycogen metabolism (Supp. Tables 6 and 7). In old mice, the proteins in glucose metabolism did not respond to the diet (Fig. 6A). In contrast, in young mice 11 proteins responded to the diet, as reflected by a deviation from the identity line in Fig. 6A, 4 of which were confirmed to be statistically significant and involved in lower glycolysis (Fig. 6B). These enzymes, namely enolase 3 (ENO3, muscle isoform), phosphoglycerate kinase 1 (PGK1), phosphoglycerate mutase 2 (PGAM2), phosphoglucomutase 1/2 (PGM1/2) and pyruvate kinase (PKM1, muscle isoform) followed a similar pattern among the tested groups: decreased by the HFD in young mice (therefore flexible to the diet) and decreased levels at old vs young age (p_age_<0.05) (Fig. 6B). Glyceraldehyde 3-phosphate dehydrogenase (GAPDH, Fig. 6B) was modulated by neither age nor diet. Besides control at the level of ATP demand, skeletal muscle glycolysis is also controlled by hexokinase (main muscle isoform in mice: HK2) (38). Age decreased HK2 content in quadriceps by about 25% (p_age_=0.026) (Fig. 6B). The proteomics results were also reflected at the level of enzyme activities. Hexokinase activity was reduced by 18.5% by advanced age (p_age_=0.03) (Fig. 6C). Pyruvate kinase activity was modulated by diet (p_diet_=0.02) and tended to be reduced by the HFD relative to the LFD, specifically in young animals (Fig. 6D). In the glycogen metabolism pathway, glycogen debranching enzyme (AGL), glycogen phosphorylase (PYGM) and phosphoglucomutase-1 (PGM) were reduced by advanced age, suggesting that glycogen breakdown could be adversely impacted by age (Supp. Table 6). Together, these results demonstrate that glucose metabolism in young mice was more flexible to changes in diet and that aged mice had a lower content of glycolytic proteins. Altogether, this is in line with reduced glucose utilization by the skeletal muscle in older animals and could further contribute to the IR phenotype.

**Figure 6:**
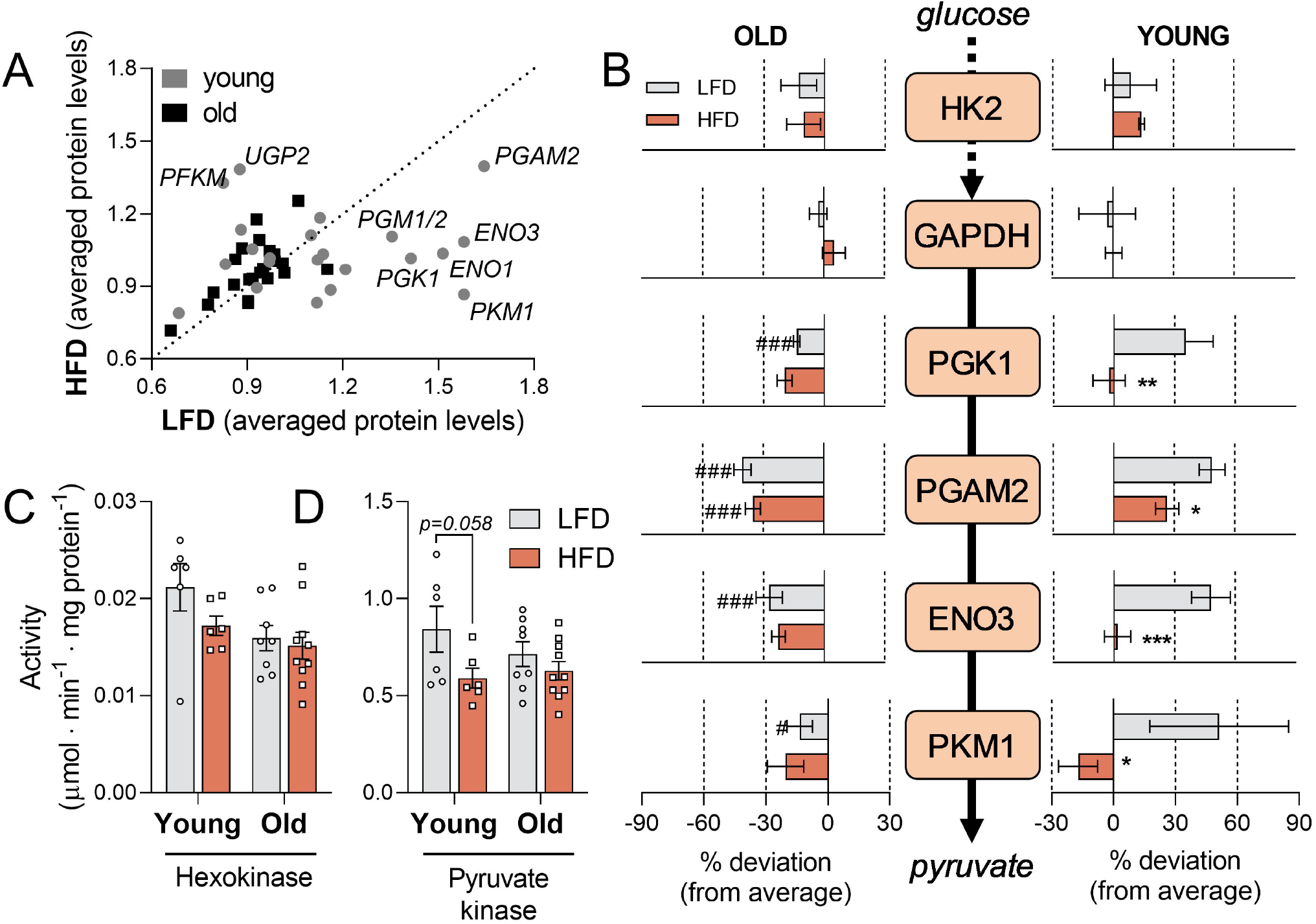
Glycolytic proteins are downregulated in aged mice, which also shows less flexibility to the diet. A: Scatter plot for proteomic targets involved in glucose metabolism (glycolysis, pentose phosphate pathway and glycogen metabolism) measured in quadriceps samples. Each dot represents the mean value of a specific protein on the HFD (y-axis) versus the LFD (x-axis). Prior to plotting, the levels of each protein were normalized to the average value of the same protein in all animals. Proteins along the dashed line are not regulated by the diet (mean LFD = mean HFD), while proteins above or below the line are up- or downregulated, respectively, by the HFD. PFKM: phosphofructokinase, muscle type, UGP2: UDP-Glucose Pyrophosphorylase 2, PGAM2; phosphoglycerate mutase 2, ENO3: enolase 3 (muscle isoform), ENO1: enolase 1, PGK1: phosphoglycerate kinase 1, PGM1/2: Phosphoglucomutase isoforms 1 and 2, PKM1: pyruvate kinase, muscle isoform. B: Simplified glycolysis scheme. The depicted values represent the % deviation from the average of the 4 groups (young/old and LFD/HFD) for each protein. The values are shown for old animals (panel on the left) and young animals (right panel). C-D: Hexokinase and pyruvate kinase activities measured in quadriceps homogenates. Data are shown as mean ± SEM and were analysed using Two-Way ANOVA and multiple comparisons were performed using Tukey’s correction. N=6-10 per group, ^###^p<0.001 (old vs young), ^##^p<0.01 (old vs young), ^#^p <0.05 (old vs young), matched diets, ***p<0.001 (LFD vs HFD), **p<0.01 (LFD vs HFD), *p<0.05 (LFD vs HFD), matched ages.

## Discussion

This study addressed the combined effects of age and diet on skeletal muscle lipid handling, and its implications for IR. We found that neither the HFD nor ageing alone led to profound lipid accumulation in the skeletal muscle, but only the combination of both. We did not observe a general decline of mitochondrial function in the quadriceps of old animals, but rather a specific failure to upregulate the β-oxidation capacity in response to the HFD, which was traced back to CPT1B. Finally, old animals had lower levels of key glycolytic proteins, which were also less responsive to the diet and could further contribute to a less flexible glucose handling.

Whereas we obtained unambiguous results concerning mitochondrial lipid handling, its relation to IR is more complex. Age strongly decreased the peripheral insulin sensitivity as quantified by the MISI index and this was further exacerbated by the HFD. The loss of peripheral insulin sensitivity in the LFD aged group, without concomitant lipid accumulation, may be explained by the progressive loss of muscle mass that accompanies ageing (16, 39). Insulin-resistant skeletal muscle displays a C18:0-ceramide signature across different species (31, 40, 41), a lipid species that directly causes IR in mice (31). Consistent with this finding, we have found a higher increase of long-chain ceramides such as C18:0 and C22:0 in old animals than in young animals in response to a HFD. Nonetheless, we failed to find a direct association between C18:0-ceramide and the MISI index. This may be related to quadriceps not being the exclusive contributor to peripheral insulin sensitivity. It may also be due, however, to modifying effects of other lipids. We found, for instance, significant negative correlations between C18:0-containing DG’s and MISI (Fig. 2M), which could indicate that C18:0-ceramide acts in synergy with DG’s to cause IR. Two of the 5 most abundant DG’s in the tissue correlated with MISI, namely DG (36:2) and DG(18:0_20:4). Interestingly, in the total membrane fraction from vastus lateralis (which belongs to the quadriceps group), DG(C18:0_20:4) also correlated negatively with insulin sensitivity in obese and diabetic individuals (42). However, a few mechanistic studies have pointed to a causal relationship between specific DG species and IR (43).

Diet and age did not have a major influence on mitochondrial biogenesis and content, as indicated by PGC-1α and citrate synthase activity, respectively. However, the proteome remodelling of young mice increased the β-oxidation capacity of quadriceps mitochondria in response to a HFD. This phenomenon appears to be a physiological response to increased lipid availability (16, 44) and may compensate for the fatty-acid overload, thereby preventing lipid accumulation (9, 13). Proteome remodelling was specific for the β-oxidation pathway and was not observed for the TCA cycle and ETC, in agreement with previous studies (35). This HFD-induced upregulation of oxidative capacity was blunted in aged animals (FC 1.8 in young vs 1.2 in old mice), despite a general upregulation of proteins involved in the β-oxidation.

The most prominent exception in the β-oxidation proteome was CPT1B, which showed decreased content and activity in old compared to young mice, and was hardly upregulated by the HFD in old mice. This observation is consistent with mRNA data from the quadriceps of aged C57BL mice (45, 46), suggesting that at least part of the defect could be at the transcriptional level. CPT1B has, in addition, been shown to be reduced by ageing in fast IIa fibres from human vastus lateralis (47) and to be decreased in obese versus lean individuals (48). In agreement with previous findings (32), we observed a shift of flux control from CPT1B at low levels of palmitoyl-CoA towards MCKAT at high levels of palmitoyl-CoA, particularly in the absence of malonyl-CoA. This may indicate that the pathway operates close to a shift of flux control, yet the integrated results indicate CPT1B as the key enzyme in the mitochondrial flexibility under the studied conditions. Therefore, an age-dependent CPT1B decline (reducing the demand for fatty acids) could be the culprit to lead to lipid accumulation when combined with increased fatty-acid supply. The latter can be caused by the HFD itself, but might be aggravated by the upregulation of CD36 (fatty-acid transport facilitator) that accompanies the HFD (20). Based on these results we propose that a reduction in skeletal muscle CPT1B with age plays a key role to predispose individuals to diet-induced insulin resistance.

Although most studies focus on the relationship between lipids and insulin signalling (1), glucose utilization in response to insulin can be partly regulated via the content of glycolytic enzymes. Upon insulin stimulation or exercise, for instance, the control of glucose utilization can shift from glucose uptake towards hexokinase (38). In the present study, we observed decreased content and flexibility of several glycolytic proteins and activities in aged animals, including enzymes that may share the control of flux, such as hexokinase and pyruvate kinase (38, 49). These results are in line with the previously reported trend of decreased levels of enzymes of lower glycolysis in old C57BL/6 mice (47, 50). Quadriceps is a fast-twitch muscle in which type II fibres predominate. We found earlier that the contribution of type IIb fast glycolytic fibres decreased with age and by a high-fat/sucrose diet, while that of type IIa fast-twitch oxidative-glycolytic fibres increased (16). The changes that we found in the proteome, at the level of both lipid and glucose metabolism, are in agreement with such modulation of fibre type.

## Conclusion

We focused on the alterations in metabolic pathways that explain the decline of flexibility in lipid and glucose handling with age. We propose CPT1B to be the main driver of the loss of flexibility to the HFD in aged mice. Future studies should elucidate the underlying transcriptional or translational mechanisms behind its regulation with age, as well as its causative role on the flexibility to the diet. Strategies that enhance muscle CPT1B activity could, nonetheless, offer the potential to attenuate lipid accumulation and consequently lipid-induced insulin resistance in the elderly.

## Authors’ contributions

MAVL, BMB, JWJ and JKK conceptualized the project, MAVL, MBD, BMB, JKK and JWJ designed experiments, MAVL, MBD, MB, WZ, JCW, AG and YvdV performed experiments, MAVL, MBD, WZ, JCW and TvD performed data analysis, RvO provided the animals used for this research, FA contributed to computer simulations and DJR contributed to lipidomics. JWJ, JKK and BMB supervised the project, MAVL and BMB wrote the manuscript, and all authors provided feedback on the manuscript.

## Acknowledgements

We thank Pim de Blaauw, Angelika Jurdzinski and Rick Havinga for their technical assistance.

## Ethics approval

All experimental procedures were approved by the institutional ethical committee of the University of Groningen

## Availability of data and materials

Large datasets generated during and/or analysed during the current study are included in the online supplementary files. Further datasets are available from the corresponding author upon reasonable request.

## Consent for publication

Not applicable.

