## Supplementary figures and images for "Age-related susceptibility to insulin resistance is due to a combination of CPT1B decline and lipid overload"

### Supplemental Figure 1

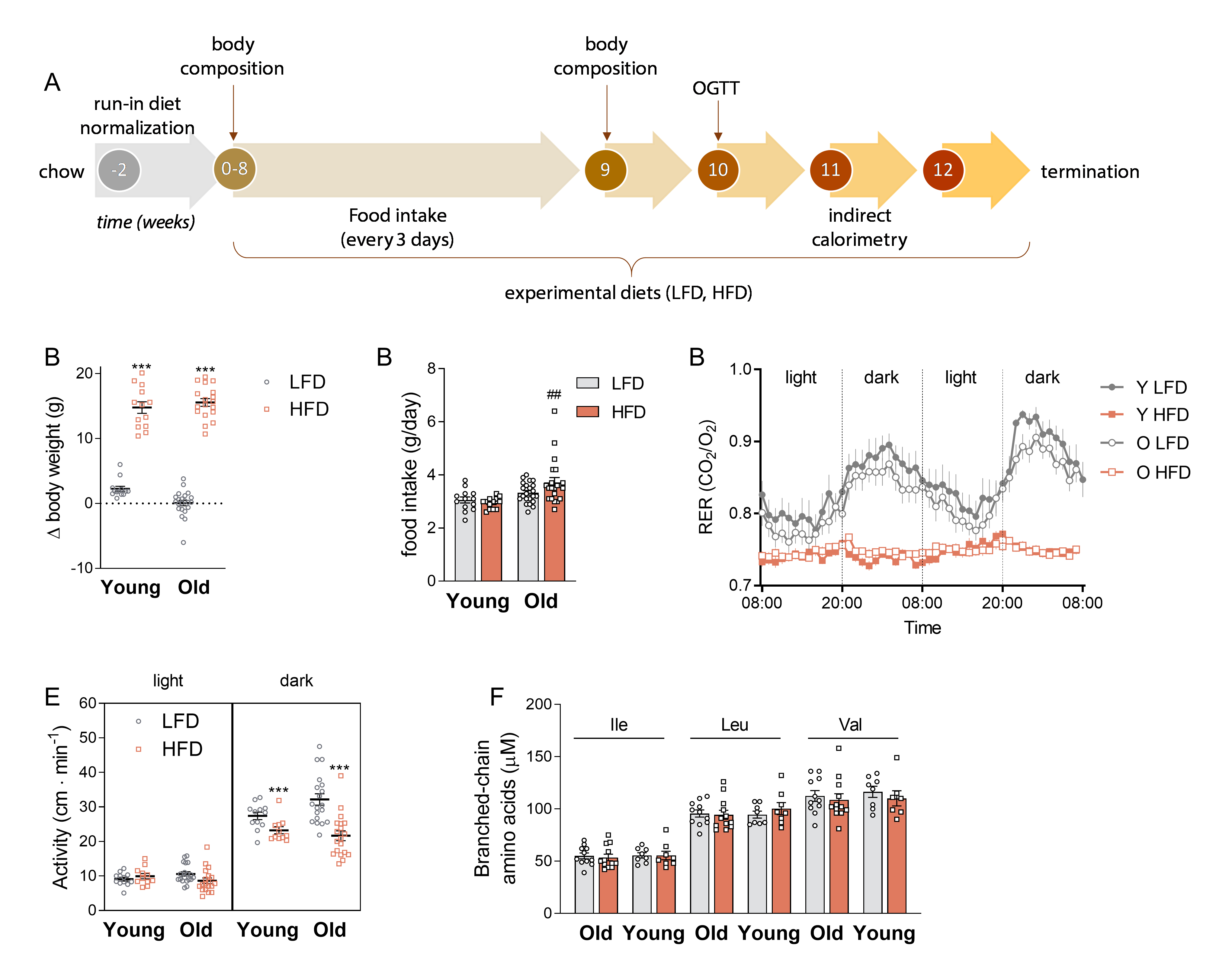

### Supplemental Figure 2

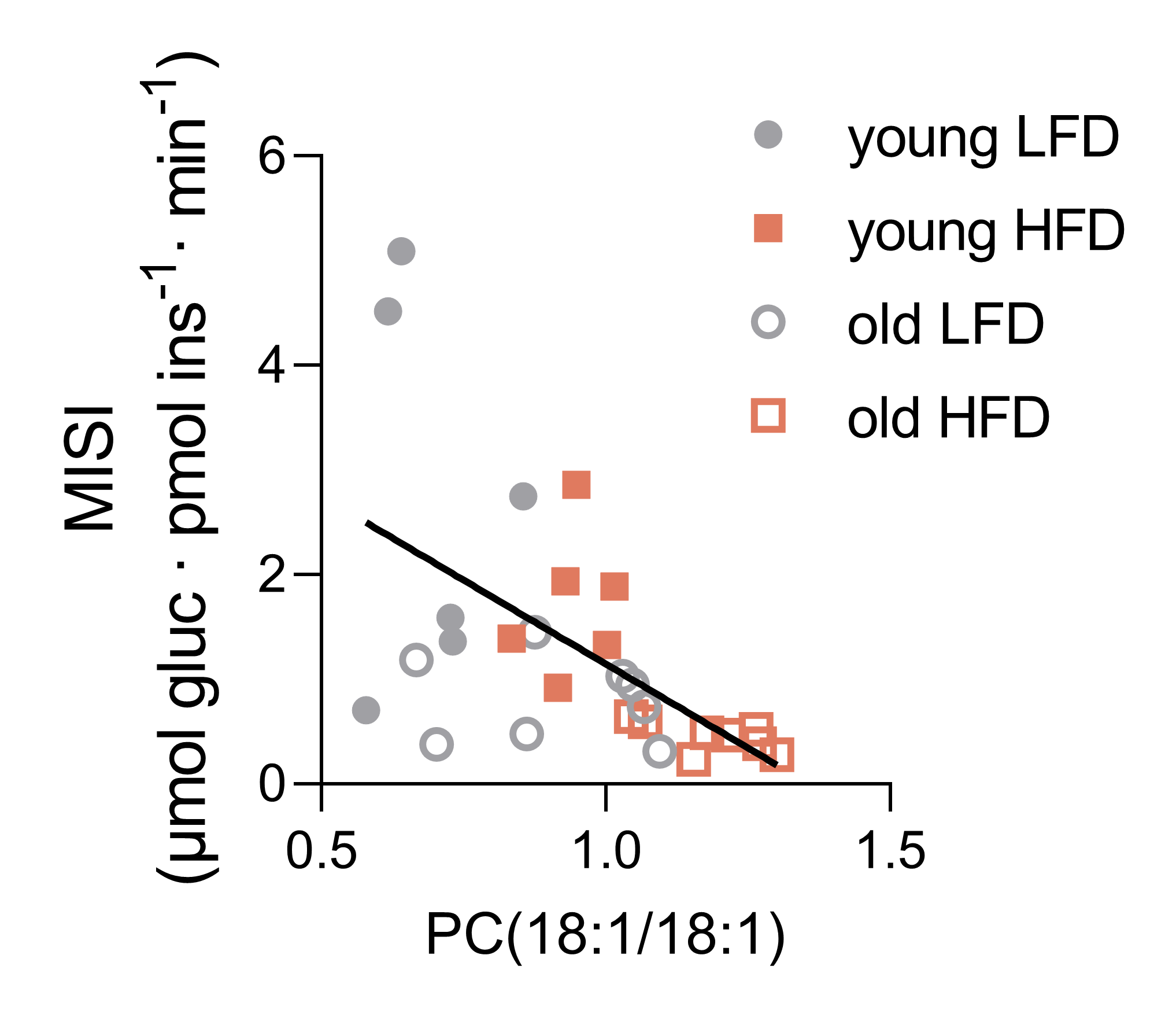

### Supplemental Figure 3

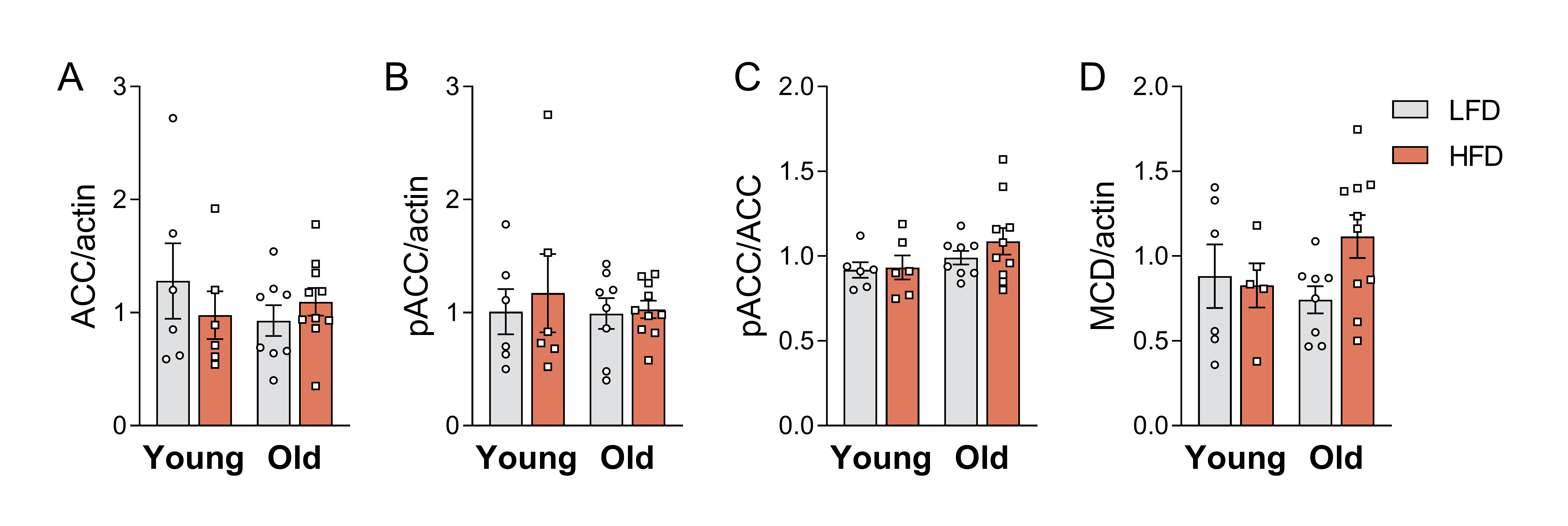
