## Supplemental Material for "Age-related susceptibility to insulin resistance is due to a combination of CPT1B decline and lipid overload"

**Mouse experiments**

The mouse experiments here reported have been part of a larger study performed at the University Medical Center Groningen, the Netherlands. Therefore, both old and young control mice on a low-fat diet (LFD) correspond to mice on a medium-protein diet (MP), as reported by Dommerholt et. al, 2020 (1).

**Physiological parameters**

In order to assign animals to either a low- or high-fat diet, old animals were normalized for body weight and 4 h fasting glucose, insulin and cholesterol levels before division into experimental groups. Young animals were randomly placed on one of the experimental diets. Animals were weighed weekly and body composition was determined, both during the run-in period and after 8 weeks of the experimental diet, using a Bruker's Minispec Whole Body Composition Analyzer. Food intake was measured several times over a 72h period. Energy expenditure (EE) and respiratory exchange ratio (RER) were measured using LabMaster metabolic cages (TSE systems, Bad Homburg, Germany). Mice were acclimatized to the cages overnight before measurements were recorded for 48 hours. The system measured O_2_ consumption and CO_2_ production to calculate the EE. The RER was determined as the ratio of the volumes of produced CO_2_ versus consumed O_2_. Infrared beams recorded locomotor activity according to the number of beam break events in the horizontal (x) and vertical (z) plane.

In terminal blood samples, plasma triglycerides, total cholesterol, and free fatty acids were determined with commercially available kits (Roche Diagnostics, Mannheim, Germany and DiaSys DiagnosticSystems, Holzheim, Germany). Amino-acid levels were quantified using cation-exchange high-performance liquid chromatography followed by post-column ninhydrin derivatization, on a Biochrom30 analyzer (Pharmacia Biotech, Cambridge, UK) (2). Norleucine was used as an internal standard.

**Assessment of glucose and insulin tolerance**

To determine changes in glucose homeostasis and insulin sensitivity, an oral glucose tolerance test (OGTT) was performed following oral administration of 1.5g/kg body weight D-glucose after overnight (10h) fasting. A 25% glucose solution (w/v) was given by oral gavage. Blood glucose levels were measured at 0, 5, 15, 30, 45, 60, 90, and 120 minutes after administration of the glucose bolus, using a OneTouch Select Plus glucose meter (Lifescan, Zug, Switzerland).

To measure insulin response by glucose administration, additional blood spots were taken at time points 0, 5, 15, 30, 60 and 120 minutes after glucose administration. Insulin was extracted from the blood spots and concentrations were determined using the rat insulin ELISA kit from Crystal Chem (Cat. 90010, Zaandam, The Netherlands) and a mouse insulin standard (Cat. 90020, Zaandam, The Netherlands) according to the manufacturer’s protocol. Homeostatic Model Assessment of Insulin Resistance (HOMA-IR) indexes were calculated as previously described using both the fasting insulin and fasting glucose levels and corrected for mice (3).

To avoid reinfusing blood cells as performed in frequently sampling protocols, the relatively small datasets of blood glucose and plasma insulin over time were fitted to equation 1 below with the use of SAAM II v2.1 (The Epsilon Group, Charlottesville, VA, USA), where C_t_ is the concentration of glucose over time. This was performed to generate enough data points for the calculation of the Muscle Insulin Sensitivity Index (MISI). This index has the units of a glucose clearance rate (dC_t_/dt) divided by the average insulin concentration during the OGTT (I), as previously defined (4). To overcome the need to fit a linear equation to the glucose clearance part of the glucose tolerance curve (4,5), we obtained the maximal dCt/dt after the glucose concentration peak using GraphPad Software Inc., version 8.0, 2018.

$$C_{t}= C_{b}+C_{1}(e^{-k_{e_{1}}\cdot t}-e^{-k_{a}\cdot t})+C_{2}\cdot e^{-k_{e_{2}}\cdot t} (1)$$

**Lipid extraction**

LC-MS grade acetonitrile (ACN), methanol (MeOH), isopropanol (IPA) and chloroform were purchased from Biosolve BV (Valkenswaard, The Netherlands). Ammonium formate, formic acid and methyl tert-butyl ether (MTBE) were purchased from Sigma Aldrich (St. Louis, MO). Various lipid standards were purchased from Avanti Polar Lipids, Inc. (Alabaster, AL). 20 µL of muscle homogenates (~3 mg muscle) were mixed with 300 µL MeOH. A defined amount of lipid standards was added to each tube followed by 10 min sonication [Composition: 2 nmol of phosphatidylcholine PC(15:0_18:0-d7), 2 nmol lysophosphatidylcholine LPC(18:1-d7), 3 nmol plasmenyl-PC(p18:0_18:1), 6 nmol phosphatidylinositol PI(15:0_18:1-d7), 4 nmol phosphatidylethanolamine PE(15:0_18:1-d9), 6 nmol lyso PE(18:1-d9), 3 nmol plasmenyl-PE(p18:0_18:1-d9), 4 nmol cardiolipin CL(14:0), 2 nmol sphingomyelin SM(18:1-d9), 3 nmol ceramide Cer(d18:1-d7/15:0), 10 nmol monoglyceride MG(18:1-d7), 2 nmol diglyceride DG(15:0_18:1-d7), 2 nmol DG(17:0_17:0-d5), 0.6 nmol triglyceride TG(15:0_18:1-d7_15:0), 0.6 nmol TG(17:0_17:1_17:0-d5) and 10 nmol cholesterol-d7. Subsequently, 1 mL MTBE was added to the mixture and kept under 25 °C on a shaker (900 rpm) for 30 min. Phase separation was induced by adding 190 µL H_2_O. The mixture was then centrifuged at 3000 g for 10 min and 850 µL of the upper phase were transferred to a new tube. The re-extraction was performed by adding 600 µL MTBE/MeOH/H_2_O (10:3:2.5, v/v/v) into the lower phase and 500 µL was collected after centrifugation to combine with the previous organic phase. The combined lipid extract solutions were dried in a vacuum centrifuge at 45 °C. Subsequently, dried lipid extracts were resuspended in 50 µL Chloroform/MeOH/ H_2_O (60:30:4.5, v/v/v) and further diluted with 150 µL IPA:ACN:H_2_O (2:1:1 v/v/v). 24 µL were injected in both positive and negative modes for LC-MS analysis. In addition, 20 µL of lipid solution taken from each sample were combined to generate a quality control pool sample and measured repeatedly throughout the measurements to monitor the technical reproducibility.

**Lipidomic LC-MS analysis**

LC-MS lipid analysis was performed on an Ultimate 3000 High-Performance UPLC coupled with a Q Exactive MS (Thermo Fischer Scientific, Darmstadt, Germany). Chromatography separation was achieved with an Acquity UPLC CSH column [1.7 μm, 100 × 2.1 mm, (Waters Corporation, Milford, MA)] under 55 °C with a flow rate of 0.4 mL/min. Mobile phase A was composed of H_2_O/ACN 40:60 (v/v), 10 mM ammonium formate and 0.1% formic acid. Mobile phase B contained ACN/IPA 10:90 (v/v) with 10 mM ammonium formate and 0.1% formic acid. The LC gradient ((6), modified) started with 40% B and raised up to 43% B at 2 min. The percentage of B raised up to 50% in the next 0.1 min and increased to 54% in the next 9.9 min. B raised to 70% in 0.1 min and increased to 99% in 5.9 min and maintained in 99% for 1 min. Subsequently, the percentage of B went back to 40% in 0.1 min and the system was equilibrated for 3.9 min before the next run started. The MS was configured under both positive and negative modes for data-dependent acquisition. A full MS scan ranging from 200-1750 m/z was acquired at resolution 70,000 FWHM (AGC target 1E6, maximum injection time 50 ms) followed by up to 8 MS/MS events with a collision energy at the resolution of 17,500 FWHM. Precursor isolation window was set to 1.5 Da with the dynamic exclusion time of 6 s. The ionization settings were as follows: capillary voltage: +3.2 kV; capillary temperature: 320 °C; sheath gas/auxiliary gas: 60/20.

**Lipidomic data processing**

Thermo Xcalibur® software [(version 3.2.63), Thermo Scientific, Waltham, MA] was used for data acquisition. Progenesis QI® software (Waters Corporation, Milford, MA) was used for data preprocessing including retention time alignment, peak picking and annotation of the LC-MS data. Peak picking was performed with an absolute ion intensity filter of 200,000 counts. Lipid annotation was performed by searching published lipid databases (Human Metabolome Database and Lipidmaps) based on mass accuracy (<5ppm), isotopic similarities, adduct type, MS/MS spectra and elution behaviour. Analysis of the fatty-acyl composition of selected lipids was performed with Lipidhunter2 software (7). Lipids intensities were normalized to the intensities of corresponding lipid standards from the same class. In case of the absence of lipid standards, the intensity was normalized to the average of all standards.

**Acylcarnitine profiling**

Acylcarnitines were measured in 15% quadriceps homogenate prepared in DPBS (Gibco 14190) using a BeadBeater system. 10 µL was collected per sample and 100 µL of acetonitrile was added, followed by the addition of 100 µL of internal standard ([8,8,8-^2^H_3_]-octanoyl-L-carnitine and [10,10,10-^2^H_3_]-decanoyl-L-carnitine). Samples were then centrifuged at 20,000 g for 10 minutes, collected into glass vials and analysed as previously described (8).

**Mitochondrial isolation and respiration**

The quadriceps was dissected on ice to remove the excess of fat, finely minced in isolation buffer (220 mM mannitol, 70 mM sucrose, 5 mM 2-[Tris(hydroxymethyl)-methylamino]-ethanesulfonic acid (TES), 0.1 mM EGTA, pH 7.3) supplemented with 0.2 mg/mL proteinase (Sigma P8038) and incubated for 5 minutes. Subsequently, BSA was added to a final concentration of 0.5 mg/mL and homogenized using a Potter system (VWR, VOS power basic). Samples were centrifuged at 800 g for 10 minutes, the pellet was discarded and the remaining supernatant was centrifuged at 7,200 g for 10 minutes. The pellet was then resuspended in 5 mL of isolation buffer and centrifuged one more time at 7,200 g for 10 minutes. The remaining pellet was resuspended in mitochondrial buffer (200 mM sucrose, 10 mM Tris, pH 7.4) and protein content was determined via the Pierce BCA Protein Assay Kit (ThermoFisher 23225).

Oxygen consumption rates were measured in MiR05 buffer (respiration buffer) containing 110 mM sucrose, 60 mM potassium lactobionate, 20 mM taurine, 20 mM HEPES, 0.5 mM EGTA, 10 mM KH_2_PO_4_, 3 mM MgCl_2_, 1 mg/mL bovine serum albumin (BSA), pH 7.1 (9) at 37 °C using a two-channel high-resolution Oroboros oxygraph-2k (Oroboros, Innsbruck, Austria). Different substrate combinations were used, namely 2 mM pyruvate and 2 mM malate (with or without 5 mM glutamate) or 25 µM palmitoyl-CoA and 2 mM L-carnitine, all in the presence of 2 mM malate. Maximal ADP-stimulated respiration (state 3) was measured in the presence of 1.5 U/mL hexokinase, 10 mM glucose and 1 mM ATP. State 4 rates were measured after addition of 1.25 µM carboxyatractyloside (CAT).

**Proteomics sample preparation**

15% (w/v) quadriceps homogenates were prepared in DPBS (Gibco 14190) with a BeadBeater system (Precellys^®^ Evolution, Bertin Technologies). Samples were centrifuged at 15,000 *g* for 5 minutes, supernatants were collected and cOmplete proteinase inhibitor cocktail was added (1:25, Merck 11836145001). Protein concentrations were measured with the Pierce BCA Protein Assay Kit (ThermoFisher 23225) Protein levels of the proteins related to the mitochondrial and glycose pathways were quantified from the skeletal muscle homogenates using targeted proteomics (10). Briefly, in-gel digestion was performed on 50 µg total protein for the skeletal muscle homogenates using trypsin (1:100 g/g sequencing grade modified trypsin V5111; Promega) after reduction with 10 mmol/L dithiothreitol and alkylation with 55 mmol/L iodoacetamide proteins, followed by solid-phase extraction (SPE C18-Aq 50 mg/1 mL, Gracepure, Thermo Fisher Scientific) for sample clean-up.

**Proteomics LC-MS analysis**

Liquid chromatography (LC) on a nano-ultra high-performance liquid chromatography (UHPLC) system (Ultimate UHPLC focused; Dionex, Thermo Fisher Scientific) was performed to separate the peptides using a nanocolumn (Acclaim PepMap100 C18, 75 µm × 500 mm 2 µm, 100 Å) with a linear gradient from 3% to 60% v/v acetonitrile plus 0.1% v/v formic acid in 110 min at a flow rate of 200 nL/min. The target peptides were analyzed by a triple quadrupole mass spectrometer (MS) equipped with a nano-electrospray ion source (TSQ Vantage; Thermo Scientific). For the LC-MS measurements, an amount of the digested peptides equivalent to a total protein amount of 1 µg total protein starting material was injected together with 0.2 (low abundant subset targets) or 1 (high abundant subset targets) ng isotopically-labelled concatemer-derived standard peptides (QconCAT technology, PolyQuant GmbH Germany, containing ^13^C-labeled arginines and lysines), plus 5 pmol GAPDH and 1 fmol SLC25a20 isotopically-labelled standard peptides (PEPotec grade 2, Thermo Scientific, containing ^13^C^15^N-labeled arginines and lysines).

**Proteomics data processing**

The MS traces were manually curated using the Skyline software (11) prior to integration of the peak areas for quantification. The sum of all transition peak areas for the endogenous peptide and isotopically labelled-peptide standard was used to calculate the ratio between the endogenous and standard peptides. The concentrations of the endogenous peptides were calculated from the known concentration of the standard and expressed in femtomoles per microgram of total protein. For proteins with more than one detected peptide, values were averaged in order to obtain an estimation for the total protein concentration. For the newly developed peptides spanning proteins in glucose metabolism (Supp Table 7), the glucose transporters (GLUT family) could not be detected endogenously, probably due to a limited extraction efficiency of plasma-membrane proteins.

**Enzyme activities**

Citrate synthase (CS), hexokinase and pyruvate kinase activities were measured in homogenates prepared in the same way as described above for proteomics assay. Citrate synthase activity was measured as previously described (12) in homogenates and mitochondrial preparations. Briefly, samples were incubated in experimental buffer containing 100 mM Tris-HCl, 5 mM triethanolamine-HCl, 0.5 mM oxaloacetate, 0.1 mM dithionitrobenzoate and 0.1% Triton-X, pH 8.1. The final protein concentration of samples was between 25 to 40 µg/mL. Reactions were initiated by addition of 0.5 mM acetyl-CoA (final concentration) and thionitrobenzoate (oxidized product) production was followed by absorbance measurements at 412 nm and 37 °C for 5 minutes. The ratio between CS activity in the homogenate and the mitochondria gives the ratio between mitochondria protein per total tissue protein, thus it reflects the enrichment of mitochondrial preparations. Consequently, this ratio can be used to express results per total tissue capacity (13).

Hexokinase and pyruvate kinase activities were carried out using NAD(P)H-linked assays at 37 °C in a Synergy H4 plate reader (BioTek™) at 340 nm for 6 min as previously described (14). Briefly, for both enzyme measurements, the assay buffer contained 100 mM Tris-HCl, 15 mM NaCl, 0.5 mM CaCl_2_, 140 mM KCl and 5 mM potassium phosphate buffer (pH 7.0). For hexokinase activity measurements, 1.2 mM NADP^+^ (Sigma N0505), 10 mM Glucose (Merck 1.083420), 1.8 U/mL glucose-6-phosphate dehydrogenase (Sigma G7877), 10.5 mM MgSO_4_ and 10 mM ATP (Sigma A2383) as start reagent were used. The buffer for pyruvate kinase measurements contained 0.15 mM NADH (Sigma N8129), 1 mM ADP (Sigma A5285), 1 mM fructose 1,6-bisphosphate (Sigma F0752), 60 U/mL L-lactate dehydrogenase (Sigma L2500) and 2 mM phosphoenolpyruvate (Sigma P7252) was used as start reagent. Four dilutions were performed per sample (0.4-4.5 mg protein/mL) to check for linearity over time and samples were further diluted 50× for pyruvate kinase and 10× for hexokinase in the experimental buffer for measurements.

Carnitine palmitoyltransferase (CPT) activity was measured using mitochondrial enriched suspensions. The experimental buffer contained 25 mM Tris-HCl, 2 mM EDTA, 150 mM KCl, 5 mM KCN, 1 mg/mL BSA, 4.5 mM reduced glutathione and 0.02 mM potassium phosphate buffer (pH 7.0). 50 µM palmitoyl-CoA (Sigma P9716) and 2 mM L-carnitine (Sigma C0158) were used as substrates and the mixture was incubated for 10 min at 37 °C. Two sample dilutions were used per animal (8 or 16 ng protein/mL) and samples were collected and quenched in acetonitrile at 0, 5 or 10 minutes (adapted from (15)). The concentration of palmitoyl-carnitine produced was measured according to the acylcarnitine profiling described above after the addition of internal standard.

**Western Blots**

For Western Blot analyses, frozen quadriceps samples were used to prepare 7.5% homogenates (w/v) in 0.1% NP-40 buffer (0.4 M NaCl) in the presence of 1 tablet of cOmplete protease inhibitor cocktail (Merck, 11836145001), 0.5 mL phosphatase inhibitor cocktail 2 (Sigma-Aldrich P5726), 0.5 mL phosphatase inhibitor cocktail 3 (Sigma-Aldrich P0044) and 1 mM dithiothreitol (DTT). Samples were homogenized in a BeadBeater, mixed with SDS loading buffer and heated at 95 °C for 5 minutes. SDS-PAGE was used to separate proteins in 8% acrylamide gels. 90 to 150V were used to separate proteins using a MiniPROTEAN Tetra Vertical Electrophoresis Cell system (Cat. No. 1658029FC; Bio-Rad) in the presence of running buffer (0.2 M glycine, 25 mM Tris and 0.1% SDS). Transfer was performed using polyvinylidene difluoride (PVDF) in blotting buffer (0.1 M glycine, 50 mM Tris, 0.01% SDS and 10% methanol, pH 8.3) at 45 V for 1 h 45 min. Membranes were blocked in 5% BSA in TBST for 1h and primary antibodies were incubated overnight at 4 °C. Subsequently, membranes were washed three times with TBST and incubated with HRP-coupled secondary antibody (goat anti-mouse or goat anti-rabbit) for 2h followed by wash before detection. Pierce ECL substrate (Thermo Fisher Scientific, 32209) was used for detection using Image Quant LAS4000 Mini (GE Healthcare) (16). Antibodies used: actin (Millipore MAB1501, 1:10,0000), ACC (Cell Signalling, 3676, 1:1,000), pACC(S79) (Cell Signalling, 3661, 1:2,000), MCD (Abcam, ab95945, 1:1,000), GAPDH (Abcam, ab37187, 1:20,000) and PGC1α (Abcam, ab54481, 1:1,000).

**Computational modelling**

The computational model of mouse mitochondrial β-oxidation previously described (17) was parameterized on an individual mouse basis. This was based on the quantitative proteomics data obtained for this study, thereby yielding a set of results for each experimental group. The rate through the CPT1 reaction with palmitoyl-CoA as substrate (vcpt1c16 in the computational model of the mitochondrial β-oxidation) was used as the β-oxidation flux in each simulation. Flux control coefficients (FCC) were calculated as previously described (18). The definition of the parameter and the estimation used for calculations can be found in equations 1 and 2, respectively. It corresponds to which extent a certain enzyme controls the flux (J) through a pathway (∆V_max_ = V_max_∙10^-6^ µmol/min^-1^∙mg protein^-1^). Summation theorem was checked (19).

$${FCC}_{enzyme}^{J}= \frac{\partial lnJ}{\partial{lnV}_{max}} \left( 1 \right)$$

$${FCC}_{enzyme}^{J}\approx\frac{\Delta J}{\Delta V_{max}}\cdot\frac{V_{max}}{J} \left( 2 \right)$$

**Statistical analysis**

Data were visualized using GraphPad Prism software (GraphPad Software Inc., version 8.0, 2018) and analysed with either the same software or IBM SPSS Statistics (IBM Corp., Version 25.0, 2017). Body weight and glucose and insulin time courses were analysed with 3-way ANOVA (age, diet and time). Delta fat mass, HOMA-IR, insulin action, plasma TG’s and NEFA’s were analysed by 2-way ANOVA (HOMA-IR and TG’s after log_2_ transformation) followed by Tukey’s multiple comparisons. Energy expenditure was adjusted for total body weight with ANCOVA according to the guidelines of the National Mouse Metabolic Phenotyping Centers. RER and adjusted energy expenditure were analysed with 3-way ANOVA (diet, age and light/dark cycle), followed by Holm-Sidak’s multiple comparisons test. Lipidomics data were transformed by mean centring and multiple Student’s t-tests were performed on data (comparing the effect of the diets on matched age groups) after log_2_ transformation. Correction for multiple comparisons used the False Discovery Rate (FDR) approach (Q-value = 5%). A significant change was defined as having a q value <0.05 and a fold change ≤ 0.66 or ≥ 1.5. Individual lipid species and PC/PE ratio were analysed with 2-way ANOVA followed by Tukey’s multiple comparison analysis. Association between MISI and lipid species was assessed by Pearson correlation coefficients. Remaining comparisons (Western Blots, enzyme activity, O_2_ consumption and protein data) were performed by 2-way ANOVA followed by Tukey’s multiple comparisons. The effect of each factor based on the 2-way ANOVA (p_age_ and p_diet_), as well as the interaction between them (p_age×diet_), was calculated for each comparison.

**Supplemental tables**

**Supplemental table 1: Diet composition (D17041407, -09)**

| Product # | D17041407 | | | | D17041409 | |
| --- | --- | --- | --- | --- | --- | --- |
|  | Low Fat (20 kcal %) | | | | High Fat (60 kcal %) | |
| Ingredient | (g) | |  | | (g) | |
| Casein | 220 | |  | | 220 | |
| Cystine | 3 | |  | | 3 | |
| Corn Starch | 368 | |  | | 67 | |
| Maltodextrin 10 | 122.2 | |  | | 22.3 | |
| Sucrose | 107.078 | |  | | 107.078 | |
| Cellulose | 50 | |  | | 50 | |
| Soybean Oil | 90 | |  | | 90 | |
| Lard | 0 | |  | | 178.4 | |
| t-butylhydroquinone | 0.014 | |  | | 0.014 | |
| Mineral Mix S10022C | 3.5 | |  | | 3.5 | |
| Calcium Carbonate | 12.495 | |  | | 12.495 | |
| Potassium Citrate, 1 H20 | 1.773 | |  | | 1.773 | |
| Potassium Phosphate, monobasic | 7.763 | |  | | 7.763 | |
| Sodium Chloride | 2.59 | |  | | 2.59 | |
| Vitamin Mix V10037 | 10 | |  | | 10 | |
| Choline Bitartrate | 2.5 | |  | | 2.5 | |
| FD&C Yellow Dye #5 | 0.05 | |  | | 0 | |
| FD&C Red Dye #40 | 0 | |  | | 0.025 | |
| FD&C Blue Dye #1 | 0 | |  | | 0.025 | |
| Total | 1000.96 | |  | | 778.463 | |
| kcal/gm | 4 | |  | | 5.2 | |
|  | gram | gram% | | gram | | gram% |
| Protein | 197 | 20 | | 197 | | 25 |
| Carbohydrate | 607 | 61 | | 206 | | 27 |
| Fat | 90 | 9 | | 268 | | 34 |
| Fiber | 50 | 5 | | 50 | | 6 |
|  | kcal | kcal% | | kcal | | kcal% |
| Protein | 786 | 20 | | 786 | | 20 |
| Carbohydrate | 2429 | 60 | | 826 | | 20 |
| Fat | 810 | 20 | | 2416 | | 60 |
| Total | 4026 | 100 | | 4028 | | 100 |
|  | gram | gram% | | gram | | gram% |
| Calcium | 5 | 0.5 | | 5 | | 0.64 |
| Phosphorus | 3.53 | 0.35 | | 3.53 | | 0.45 |
| Potassium | 3.6 | 0.36 | | 3.6 | | 0.46 |
|  | Typical Amino Acids (g) | | |  | | |
| *Essential Amino Acids* | per kg diet | per 1000 kcal | | per kg diet | | per 1000 kcal |
| Histidine | 4.95 | 1.23 | | 6.36 | | 1.23 |
| Isoleucine | 8.24 | 2.05 | | 10.6 | | 2.05 |
| Leucine | 17.25 | 4.29 | | 22.18 | | 4.29 |
| Lysine | 14.29 | 3.55 | | 18.37 | | 3.55 |
| Methionine | 5.49 | 1.37 | | 7.07 | | 1.37 |
| Phenylalanine | 9.12 | 2.27 | | 11.73 | | 2.27 |
| Threonine | 7.8 | 1.94 | | 10.03 | | 1.94 |
| Tryptophan | 2.31 | 0.57 | | 2.97 | | 0.57 |
| Valine | 10.11 | 2.51 | | 13 | | 2.51 |
| *Non-Essential Amino Acids* |  |  | |  | |  |
| Alanine | 5.49 | 1.37 | | 7.07 | | 1.37 |
| Arginine | 6.48 | 1.61 | | 8.34 | | 1.61 |
| Asparagine | 7.69 | 1.91 | | 9.89 | | 1.91 |
| Aspartate | 5.49 | 1.37 | | 7.07 | | 1.37 |
| Cystine | 4.32 | 1.07 | | 4.7 | | 1.07 |
| Glutamine | 18.68 | 4.65 | | 24.02 | | 4.64 |
| Glutamate | 22.75 | 5.66 | | 29.25 | | 5.65 |
| Glycine | 3.3 | 0.82 | | 4.24 | | 0.82 |
| Proline | 19.34 | 4.81 | | 24.87 | | 4.81 |
| Serine | 10.88 | 2.71 | | 13.99 | | 2.7 |
| Tyrosine | 9.89 | 2.46 | | 12.72 | | 2.46 |

**Supplemental Table 2: Attached Excel file**

**Supplemental table 3: Lipids that are changed by a HFD in either old or young mice.**

| **Lipid name** | **Detailed composition** | **Significant result LFD vs HFD** | **Fold change** | **q-value** |
| --- | --- | --- | --- | --- |
| **TG(54:1)** | **TG(18:0_18:0_18:1)** | old | 7.7 | 0.0017 |
| **TG(41:1)** |  | old | 6.0 | 0.0258 |
| **23OH-C20:0-carnitine** |  | old | 5.7 | 0.0035 |
| **TG(52:0)** | **TG(16:0_18:0_18:0)** | old | 5.5 | 0.0017 |
| **C18:0-carnitine** |  | old | 4.9 | 0.0005 |
| **DG(36:1)** | **DG(18:0_18:1)** | old | 4.3 | 0.0025 |
| **TG(50:0)** | **TG(16:0_16:0_18:0)** | old | 4.1 | 0.0022 |
| **TG(53:1)** | **TG(17:0_18:0_18:1)** | old | 3.9 | 0.0137 |
| **C16:0-carnitine** |  | old | 3.8 | 0.0055 |
| **CL(72:8)** | **CL(18:2/18:2/18:2/18:2)** | old | 3.5 | 0.0129 |
| **TG(54:2)** | **TG(18:0_18:1_18:1)** | old | 3.4 | 0.0147 |
| **3OH-C18:1-carnitine** |  | old | 3.2 | 0.0297 |
| **TG(51:1)** | **TG(16:0_17:0_18:1)** | old | 2.9 | 0.0317 |
| **PC(40:4)** | **PC(18:0_22:4)** | old | 2.8 | 0.0033 |
| **TG(51:0)** | **TG(16:0_17:0_18:0)** | old | 2.7 | 0.0156 |
| **TG(59:2)** |  | old | 2.7 | 0.0499 |
| **DG(38:4)** | **DG(16:0_22:4)** | old | 2.7 | 0.0054 |
| **C14-carnitine** |  | old | 2.6 | 0.0352 |
| **PC(38:4)** | **PE(16:0_22:4)** | old | 2.6 | 0.0352 |
| **TG(42:0)** | **TG(12:0_14:0_16:0)** | old | 2.3 | 0.0240 |
| **PC(37:4)** | **PC(17:0_20:4)** | old | 2.2 | 0.0005 |
| **PC(37:3)** |  | old | 2.2 | 0.0189 |
| **PEp(40:5)/PEa(40:6)** |  | young/old | 2.2/1.9 | 0.0438/0.0129 |
| **PEa(40:5)/PEp(40:4)** |  | young/old | 2.1/2.2 | 0.0307/0.0021 |
| **SM(d36:2)** |  | young/old | 2.1/2.2 | 0.0264/0.0005 |
| **TG(49:0)** | **TG(16:0_16:0_17:0)** | old | 2.1 | 0.0475 |
| **SM(d34:2)** |  | old | 2.1 | 0.0119 |
| **PC(38:2)** |  | old | 2.1 | 0.0033 |
| **PC(38:3)** | **PC(18:0_20:3)** | old | 2.1 | 0.0116 |
| **DG(32:0)** | **DG(16:0/16:0)** | old | 2.0 | 0.0035 |
| **PC(39:5)** |  | young/old | 1.9/2.0 | 0.0198/0.0033 |
| **Cer(d36:1)** | **Cer(d18:1/18:0)** | old | 1.9 | 0.0033 |
| **DG(34:0)** | **DG(16:0_18:0)** | old | 1.9 | 0.0005 |
| **DG(35:2)** |  | old | 1.9 | 0.0309 |
| **PC(38:4)** | **PC(16:0_22:4)** | young/old | 1.8/2.6 | 0.0198/0.0001 |
| **PC(36:2)** | **PC(18:0_18:2)** | old | 1.8 | 0.0071 |
| **PC(35:2)** | **PC(17:0_18:2)** | old | 1.8 | 0.0021 |
| **PCa(38:4)/PCp(38:3)** |  | old | 1.8 | 0.0129 |
| **PC(38:4)** | **PC(18:0_20:4)** | old | 1.8 | 0.0035 |
| **PC(36:3)** | **PC(16:0_20:3)** | old | 1.8 | 0.0055 |
| **LPE(18:0)** |  | old | 1.8 | 0.0054 |
| **LPC(20:4)** |  | old | 1.8 | 0.0147 |
| **SM(d40:2)** |  | old | 1.8 | 0.0317 |
| **LPC(22:5)** |  | old | 1.7 | 0.0335 |
| **Cer(d40:1)** | **Cer(d18:1/22:0)** | old | 1.7 | 0.0156 |
| **LPC(18:0)** |  | old | 1.7 | 0.0182 |
| **PC(33:0)** |  | old | 1.7 | 0.0116 |
| **PC(36:4)** | **PC(16:0_20:4)** | old | 1.7 | 0.0054 |
| **PC(35:1)** |  | young/old | 1.6/1.8 | 0.0198/0.0035 |
| **PC(38:5)** | **PC(16:0_22:5) major** | old | 1.6 | 0.0220 |
| **PEp(38:4)/PEa(38:5)** |  | old | 1.6 | 0.0054 |
| **PEp(38:4)/PEa(38:5)** |  | old | 1.6 | 0.0303 |
| **PC(36:1)** | **PC(18:0_18:1)** | old | 1.6 | 0.0198 |
| **DG(38:6)** | **DG(16:0_22:6)** | old | 1.6 | 0.0208 |
| **PEp(38:5(OH))** |  | young/old | 1.5/1.7 | 0.0307/0.0035 |
| **SM(d35:1)** |  | old | 1.5 | 0.0116 |
| **DG(38:4)** | **DG(18:0_20:4)** | old | 1.5 | 0.0261 |
| **PC(34:2)** | **PC(16:0_18:2) major** | old | 1.5 | 0.0129 |
| **DG(33:2)** |  | old | 0.6 | 0.0137 |
| **PC(32:2)** | **PC(14:0_18:2)** | young | 0.5 | 0.0209 |
| **PC(34:4)** |  | young/old | 0.4/0.6 | 0.0198/0.0147 |
| **TG(59: 3)** |  | old | 0.4 | 0.0178 |

**Supplemental table 4: Acylcarnitine composition in quadriceps (nmol * g tissue ^-1^)**

|  | YOUNG | | | | OLD | | | |
| --- | --- | --- | --- | --- | --- | --- | --- | --- |
|  | **LFD** | | **HFD** | | **LFD** | | **HFD** | |
| Species | **Mean** | **SEM** | **Mean** | **SEM** | **Mean** | **SEM** | **Mean** | **SEM** |
| total | 129.8 | 9.4 | 145.7 | 3.6 | 165.7 | 20.1 | 162.6 | 6.6 |
| C0 | 56.5 | 4.2 | 67.6 | 2.3 | 71.4 | 8.5 | 73.0 | 3.4 |
| C2* | 57.0 | 5.2 | 63.8 | 4.4 | 81.1 | 10.7 | 72.4 | 4.7 |
| C3 | 0.47 | 0.05 | 0.49 | 0.04 | 0.59 | 0.08 | 0.57 | 0.04 |
| C4 | 0.85 | 0.11 | 0.86 | 0.05 | 0.89 | 0.11 | 1.02 | 0.06 |
| C5 | 0.12 | 0.01 | 0.16 | 0.01 | 0.16 | 0.02 | 0.23 | 0.05 |
| C6 | 0.24 | 0.03 | 0.29 | 0.01 | 0.25 | 0.03 | 0.31 | 0.01 |
| C8 | 0.14 | 0.01 | 0.16 | 0.02 | 0.13 | 0.01 | 0.15 | 0.01 |
| C10:1 | 0.03 | 0.01 | 0.02 | 0.00 | 0.02 | 0.01 | 0.02 | 0.00 |
| C10 | 0.05 | 0.01 | 0.04 | 0.01 | 0.03 | 0.00 | 0.04 | 0.01 |
| C12:1 | 0.03 | 0.01 | 0.02 | 0.00 | 0.02 | 0.01 | 0.02 | 0.00 |
| C12 | 0.09 | 0.02 | 0.05 | 0.02 | 0.05 | 0.01 | 0.06 | 0.01 |
| C14:1 | 0.28 | 0.07 | 0.17 | 0.04 | 0.21 | 0.04 | 0.22 | 0.05 |
| C14 | 0.83 | 0.14 | 0.69 | 0.16 | 0.52 | 0.09 | 0.69 | 0.12 |
| C16:1 | 0.84 | 0.16 | 0.47 | 0.11 | 0.66 | 0.14 | 0.59 | 0.14 |
| C16 | 3.92 | 0.70 | 3.51 | 0.97 | 2.35 | 0.46 | 4.00 | 0.73 |
| C18:2 | 1.05 | 0.23 | 0.54 | 0.13 | 0.83 | 0.18 | 0.81 | 0.21 |
| C18:1 | 1.92 | 0.36 | 1.46 | 0.38 | 1.83 | 0.35 | 2.33 | 0.66 |
| C18 | 1.03 | 0.18 | 1.22 | 0.30 | 0.78 | 0.15 | **1.49^#^** | 0.23 |
| C12OH | 0.07 | 0.01 | 0.10 | 0.01 | 0.10 | 0.04 | 0.14 | 0.02 |
| C14OH | 0.08 | 0.01 | 0.10 | 0.01 | 0.06 | 0.01 | 0.10 | 0.01 |
| C16OH^$^ | 0.26 | 0.03 | 0.35 | 0.05 | 0.19 | 0.03 | **0.36^&^** | 0.03 |
| C18:1OH^$^ | 0.25 | 0.04 | 0.34 | 0.05 | 0.26 | 0.04 | **0.43^#^** | 0.05 |
| C18OH^$^ | 0.09 | 0.01 | 0.14 | 0.03 | 0.08 | 0.01 | **0.17^+^** | 0.01 |

*p<0.05 for age and $p<0.01 for diet, 2-way ANOVA. #p<0.05 for LFD vs HFD, old, &p<0.01 for LFD vs HFD, old, and +p<0.001 for LFD vs HFD, old.

**Supplemental Table 5: Correlations (Pearson r) between peripheral insulin sensitivity and ceramides/diacylglycerol levels**

| Species | Lipid | Pearson r | p value |
| --- | --- | --- | --- |
| Cer01 | Cer(d36:1) | -0.20 | 0.308 |
| Cer02 | Cer(d38:3) | -0.26 | 0.173 |
| Cer03 | Cer(d40:1) | -0.30 | 0.117 |
| Cer04 | Cer(d42:2) | -0.30 | 0.125 |
| Cer05 | Cer(d42:1) | -0.26 | 0.176 |
| Cer06 | Cer(d34:1) | -0.27 | 0.172 |
| Cer07 | Cer(d30:1) | 0.00 | 0.989 |
| Cer08 | Cer(d35:0) | -0.28 | 0.149 |
| Cer09 | 1-O-eicosanoyl-Cer(d34:1) | -0.27 | 0.168 |
| Cer10 | PE-Cer(36:3) | 0.14 | 0.486 |
| Cer11 | HexCer(d32:1) | 0.16 | 0.406 |
| Cer12 | HexCer(d38:1) | 0.25 | 0.207 |
| Cer13* | HexCer(d41:1) | -0.33 | 0.089 |
| Cer14 | HexCer(d40:1(2OH)) | -0.24 | 0.215 |
| Cer15 | HexCer(d42:1(2OH)) | -0.24 | 0.220 |
| Cer16 | HexCer(d42:1) | -0.27 | 0.158 |
| Cer17 | HexCer(d34:1) | 0.21 | 0.278 |
| Cer18 | NeuAcα2-3Galβ1-4Glcβ-Cer(d36:1) | 0.23 | 0.247 |
| Cer19 | GlcNAcβ1-3(GalNAcβ1-4)Galβ1-4Glcβ-Cer(d36:1) | -0.03 | 0.860 |
| DG01 | DG(32:0) | -0.02 | 0.914 |
| DG02 | DG(32:1) | -0.10 | 0.624 |
| DG03 | DG(32:1) | -0.02 | 0.937 |
| DG04 | DG(32:2) | 0.04 | 0.831 |
| DG05 | DG(32:2) | 0.00 | 0.981 |
| DG06 | DG(33:2) | 0.04 | 0.848 |
| DG07 | DG(34:0) | -0.24 | 0.228 |
| DG08* | DG(34:1) | -0.37 | 0.053 |
| DG09 | DG(34:2) | -0.26 | 0.187 |
| DG10 | DG(34:2) | -0.20 | 0.318 |
| DG11 | DG(34:3) | -0.03 | 0.893 |
| DG12 | DG(34:3) | -0.05 | 0.803 |
| DG13 | DG(35:2) | -0.29 | 0.138 |
| DG14 | DG(36:1) | -0.34 | 0.078 |
| DG15* | DG(36:2) | -0.46 | 0.015 |
| DG16 | DG(36:3) | -0.34 | 0.074 |
| DG17 | DG(36:4) | -0.06 | 0.753 |
| DG18* | DG(37:2) | -0.48 | 0.011 |
| DG19 | DG(38:2) | -0.31 | 0.114 |
| DG20 | DG(38:4) | -0.27 | 0.159 |
| DG21* | DG(38:4) | -0.45 | 0.016 |
| DG22 | DG(38:6) | -0.07 | 0.738 |
| DG23 | DG(40:6) | -0.02 | 0.902 |
| DG24 | DG(42:5) | 0.07 | 0.717 |

***p<0.05**

**Supplemental Table 6/7: Attached Excel files**

**Supplemental Figure - Legends**

**Supplemental Figure 1:** A: Schematic design of animal experiments. B: Change in body weight for both age groups after 8 weeks on either a LFD or HFD. C: Average food intake per day from several measurements over the course of 8 weeks. D: 48h time course for respiratory exchange ratios (RER) in both light/dark phases following 10 weeks for each age/diet group. E: Activity of animals (cm · min^-1^) measured in metabolic cages with the use of lasers in parallel with indirect calorimetry. F: Plasma branched-chain amino acids measured at termination blood spots for all age/diet groups. Data are shown as mean ± SEM. n=13-22 per group, ***p<0.001 (LFD vs HFD, matched age), Two-Way-ANOVA followed by Tukey’s post hoc test.

**Supplemental figure 2:** Linear regression between phosphatidylcholine PC(18:1/18:1) and MISI (Pearson r = -0.56, p = 0.002) .

**Supplemental figure 3:** A-D: Total ACC, pACC (S79), pACC/ACC and MCD levels in quadriceps homogenates normalized by actin levels.
