## Supplemental Text 1 for "Age-related susceptibility to insulin resistance is due to a combination of CPT1B decline and lipid overload"

**Supplemental Text 1: Modelling strategy and assumptions.**

The computational model of mouse liver mitochondrial β-oxidation was used as starting point [1]. Previously estimated values for K_m_’s, K_eq_’s as well as concentrations of intracellular metabolites were kept according to the liver model given that most isozymes are identical in both tissues. A step-by-step guide to the parameterization of V_max_’s to skeletal muscle are as follows:

1. Proteomics data measured in total quadriceps homogenate were converted using the citrate synthase ratio to be expressed per mitochondria protein.
2. Data were compared to a previous dataset of mitochondrial targets measured in mitochondria-enriched samples (quadriceps of 6-month old C57Bl/6J mice) [2]. This showed that the mild harvest conditions used for homogenates (necessary for enzyme kinetics measurements) had underestimated the absolute quantities of membrane-associated proteins for all four experimental groups. Therefore, based on the protein averages of the previous dataset and the here obtained data on the interplay between age and diet, we calculated ratios to correct for membrane loss and they were applied equally for all 32 mice in the dataset. Proteins that were underestimated in our data set with a fold change of at least 2 were corrected (Acaa2, Acadvl, Cpt1b, Cpt2, Hadha/Hadhb) according to Table 1A.
3. The previous publication from our group was used for the calculation of k_cat_’s (turnover ratios) for the proteins involved in β-oxidation based on the equation V_max_ = k_cat_ ∙ [E]. V_max_ represents the maximal capacity of a determined enzyme expressed in µmol∙min^-1^∙mg protein^-1^ and [E] is the absolute protein concentration expressed in pmol∙mg protein^-1^. A table containing the calculated values is shown below (Table 1B).
4. Missing enzymes on proteomics datasets:
   a. The study by van Eunen *et al*. did not include measurements of the CACT protein (Slc25a20), therefore k_cat_’s for this enzyme could not be estimated. Moreover, the peptide used for the detection of this protein is synthetic and consequently cannot be used for absolute but only for relative quantification. Therefore the original values of the V_max_ for this enzyme were kept identical to those used in the liver model. They were, however, multiplied by a factor proportional to the abundance of the Slc25a20 detected in the current study.
   b. The protein LCAD was not detected in our samples given its early elution during the liquid chromatography. Therefore, we estimated the values for its V_max_ based on the assumptions that follow. First, in both liver [1] and skeletal muscle [2] datasets, protein levels of LCAD were approximately 2 times higher than the levels of VLCAD. Second, the k_cat_ estimation on table 1B shows a value approximately 3.5 times higher for LCAD than that for VLCAD. Considering these two observations, we estimated LCAD V_max_’s for each mouse as 7 times the V_max_ calculated for VLCAD (after correction for membrane proteins).
5. Different isoforms: k_cat_’s estimated for CPT1 in liver regard the activity of mostly CPT1A, the most abundant CPT1 isoform in that tissue. Therefore, we estimated the k_cat_ for CPT1B based on our own proteomics and activity measurements. Figure 1A shows a plot of the calculated k_cat_ in skeletal muscle.
6. Besides the difference in V_max,_ CPT1B is known to possess lower affinity to L-carnitine than CPT1A. The K_m­_ used in the simulations was equal to 500 µM, obtained from measurements in human and rat skeletal muscle [3]
7. CPT1B can bind malonyl-CoA with a higher affinity than CPT1A [4]. Therefore the K_i_ used for this endogenous inhibitor was equal to 0.2 µM [5].
8. Malonyl-CoA was set as 0.2 µM (same as K_i_) for simulations were indicated. The value here used is below the expected concentrations found *in vivo* [3], which exceed the K_i_ and would lead to inhibition of β-oxidation to a very high extent. Therefore, it is hypothesized that either part of the malonyl-CoA is not directly available to CPT1 [4] or that other metabolites can compete for the binding pocket, such as acetyl-CoA [5].
9. The table containing the estimated V_max_’s can be found on supplemental table 8.
10. The rate through CPT1 when using palmitoyl-CoA as substrate(vcpt1c16) was adopted as the β-oxidation flux in each simulation.
11. The Mathematica scripts for the performed simulation can be found in Appendix 1.

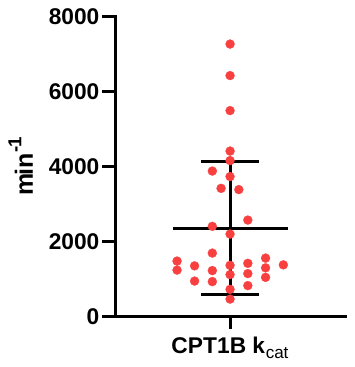

Figure 1A: Estimated k_cat_ for CPT1B (2300 ± 300 min^-1^)

Table 1A: Absolute quantification of skeletal muscle proteins corrected for mitochondrial protein. Ratios in bold represent proteins with underestimated levels in the current dataset and were used equally across all mouse groups to correct for membrane protein underestimation.

| **Enzyme (gene name)** | **Previous dataset [2]** | | **Current data (this article)** | **Ratio** | **Protein name** |
| --- | --- | --- | --- | --- | --- |
| Acaa2 | | 33.1 | 14.42 | **2.30** | 3-ketoacyl-CoA thiolase, mitochondrial (MCKAT) |
| Acadl* | | 86.9 | - | - | Long-chain specific acyl-CoA dehydrogenase, mitochondrial (LCAD) |
| Acadm | | 48.1 | 44.34 | 1.09 | Medium-chain specific acyl-CoA dehydrogenase, mitochondrial (MCAD) |
| Acads | | 17.7 | 19.39 | 0.91 | Short-chain specific acyl-CoA dehydrogenase, mitochondrial (SCAD) |
| Acadvl | | 42.3 | 4.12 | **10.27** | Very long-chain specific acyl-CoA dehydrogenase, mitochondrial (VLCAD) |
| Cpt1b | | 38.4 | 17.51 | 2.19 | Carnitine O-palmitoyltransferase 1, muscle isoform (CPT1B) |
| Cpt2 | | 16.9 | 0.60 | **27.91** | Carnitine O-palmitoyltransferase 2 (CPT2) |
| Echs1 | | 25.5 | 21.60 | 1.18 | Enoyl-CoA hydratase, mitochondrial (CROT) |
| Hadh | | 43.5 | 49.72 | 0.88 | Hydroxyacyl-coenzyme A dehydrogenase, mitochondrial (MSCHAD) |
| Hadha^$^ | | 40.3 | 6.29 | **6.42** | Trifunctional enzyme subunit alpha, mitochondrial (MTPα) |
| Hadhb^$^ | | 40.8 | 11.34 | **3.60** | Trifunctional enzyme subunit beta, mitochondrial (MTPβ) |
| Slc25a20 | | 4.4 | 1.89 | **2.33** | Mitochondrial carnitine/acylcarnitine carrier protein (CACT) |

Average in (pmol∙mg protein^-1^)

*Interference in current dataset, no peptides detected

^$^Average ratio used for MTP (Hadha/Hadhb) was 5.01

Table 1B: k_cat_ estimation based on the study by van Eunen et al. [1]

| **Enzyme (gene name)** | **Model parameter** | **V_max_** | **Protein ([E])** | **Estimated k_cat_ (min^-1^)** |
| --- | --- | --- | --- | --- |
| Acaa2 | **Vmckat** | 0.25 | 141.26 | 1,770 |
| Acadl | **Vlcad** | 0.047 | 25.45 | 1,850 |
| Acadm | **Vmcad** | 0.027 | 23.37 | 1,160 |
| Acads | **Vscad** | 0.036 | 24.72 | 1,460 |
| Acadvl | **Vvlcad** | 0.007 | 13.50 | 520 |
| Cpt2 | **Vcpt2** | 0.391 | 11.11 | 35,200 |
| Echs1 | **Vcrot** | 3.6 | 24.82 | 145,000 |
| Hadh | **Vmcschad** | 0.34 | 41.29 | 8,200 |
| Hadha/Hadhb | **Vmtp** | 2.84 | 33.49 | 85,000 |

V_max_ values in µmol∙min^-1^∙mg protein^-1^

[E] values in (pmol∙mg protein^-1^)

Values of k_cat_ expressed in s^-1^: (Vmax ∙ 10^6^)/[E]. The 10^6^ term used to convert from µmol to pmol.
